# Intronic heterochromatin prevents cryptic transcription initiation in Arabidopsis

**DOI:** 10.1101/610832

**Authors:** Jincong Zhou, Liangyu Liu, Qin Li, Wei Xu, Kuan Li, Zhi-Wei Wang, Qianwen Sun

**Affiliations:** Tsinghua-Peking Joint Center for Life Sciences and Center for Plant Biology, School of Life Sciences, Tsinghua University, Beijing 100084, China; Key Laboratory of Plant Gene Resources and Biotechnology for Carbon Reduction and Environmental Improvement, Beijing Municipal Government, and College of Life Sciences, Capital Normal University, Beijing 100048, China; Hainan Key Laboratory for Sustainable Utilization of Tropical Bioresources, College of Tropical Agriculture and Forestry, Hainan University, Haikou 570228, China

## Abstract

Intronic transposable elements (TEs) comprise a large proportion in eukaryotic genomes, but how they regulate the host genes remains to be explored. Our forward genetic screen disclosed the plant specific RNA polymerases IV and V in suppressing intronic TE-mediated cryptic transcription initiation of a chimeric transcripts at *FLC* (*FLC*^*TE*^). Initiation of *FLC*^*TE*^ transcription is blocked by the locally formed intronic heterochromatin, which is directly associated with RNA Pol V to inhibit the entry of RNA Pol II and the occupancy of H3K4 methylation. Genome-wide Pol II Ser5p native elongation transcription sequencing revealed that this is a common mechanism among intronic heterochromatin-containing genes. This study sheds light on deeply understanding the function of intronic heterochromatin on host genes expression in eukaryotic genome.

## INTRODUCTION

Many eukaryotic genomes incorporate transposable elements (TEs), which have essential roles in chromatin structure maintenance and transcriptional regulation of nearby genes in a location-dependent manner [1,2]. Moreover, functional analysis of numerous TEs inserted into intragenic regions indicates they influence multiple aspects of transcriptional regulation of their host genes, such as modulating alternative splicing and transcriptional termination [1-3]. A growing number of examples about intronic TEs play critical roles in regulating host gene expression and organism development has been reported. In the human genome, one of the largest retrotransposon families SINEs (short interspersed elements, such as Alu repetitive elements) modulates alternative splicing and induces dysfunction or neofunctionalization of corresponding proteins [4-6]. Intronic LINEs (long interspersed elements) regulate tissue-specific exon usages through recruiting RNA binding proteins to introns [7]. An intronic TE which increases host gene *Cortex* expression in the peppered moth, leading to the color adaptation to the industrial revolution [8]. A large number of transcripts identified in a cell or tissue specific manner are synthesized by TE-derived promoters with Pol II transcription initiation in human and mouse [9-11], though it is largely unclear how to specifically drive intronic TEs to be the potential transcription initiation sites.

In plants, numerous TEs are located in introns and play critical roles in gene regulation and plant development [12-16]. In Arabidopsis, the intronic TE was found to regulate the proper polyadenylation for terminating a H3K9 demethylase gene *IBM1* [17-20]. In the Arabidopsis accession Landsberg *erecta* (*Ler*), a nonautonomous Mutator-like TE had been found within the first intron of *FLOWERING LOCUS C* (*FLC*), which weakens the *FLC* function and accelerates flowering [21,22]. Interestingly, most of the Arabidopsis accessions, including Col-0, do not carry this TE insertion in the *FLC* locus [21]. In *Ler*, the mutations of HEN1 and AGO4, two essential components in small RNA biogenesis and transcriptional gene silencing [23,24], were shown to influence TE-mediated aberrant transcription on *FLC* [25], however, the detailed regulatory mechanisms are still not clear.

In this study, starting with a forward genetic screen to identify novel regulators on Mutator-containing *FLC* in *Ler* ecotype, we uncovered this intronic transposon-formed heterochromatin suppresses the cryptic transcription initiation in *FLC* intron 1. The plant specific RNA-directed DNA methylation (RdDM) pathway promotes heterochromatin formation and primarily inhibits RNA Pol II transcription initiation and active histone mark H3K4me3 switch on *FLC* as well as other targets genome-widely.

## RESULTS

### Forward screen identified plant specific RNA polymerases carrying aberrant *FLC*^*TE*^ transcripts

To investigate the regulatory mechanisms by which intronic TEs mediate aberrant RNA processing on *Ler FLC*, we developed an RT-qPCR (quantitative reverse transcription polymerase chain reaction)-based forward genetic screen (Fig S1A and S1B, Materials and Methods). We designed a primer pair based on the sequence inside the Mutator transposon and measured aberrant *FLC* transcripts (*FLC*^*TE*^, also see below) levels by RT-qPCR in an EMS-mutagenized *Ler* mutant library, with Col-0, *Ler* and *fca* mutant (which upregulates the expression of *FLC*) as negative controls (Fig S1A and S1B).

From this screen, we identified two mutants (named as *mutant a* and *b*, respectively) with significant elevated *FLC*^*TE*^ transcript levels (Fig S1B). *Mutant a* was fine-mapped to the *NRPE1* gene locus (AT2G40030) and carried a point mutation which happens on the conserved glutamic acid (Glu, E) residue (Fig S1G-J and S2A), and it is localized inside the linker region connecting NRPE1 protein domains A and B [23]. NRPE1 is the largest subunit in the plant-specific RNA Polymerase V (RNA Pol V) complex and is required for transcribing long non-coding RNA transcripts associated with AGO4-siRNA complex in the RdDM pathway [24,26]. Fine-mapping of the *mutant b* revealed it creates a premature stop codon in the thirteenth exon of *NRPD1* (AT1G63020, Fig S1C-F), which is the largest subunit of plant-specific RNA Pol IV and mediates siRNA biogenesis in the RdDM pathway [23,24]. Hereafter, we named *mutant a* and *b* as new mutations of *nrpe1-L1* and *nrpd1-L1*, respectively (Fig S1B). Consistently, results of RT-qPCR screen can be comfirmed by both Northern blot and RT-PCR (Fig S2B and S2C, Materials and Methods).

The aberrant *FLC*^*TE*^ transcript defect in *nrpe1-L1* can be complemented by *NRPE1:FLAG* driven by its native promoter (Fig S2C) [27,28]. The repressed Pol V target *AtSN1* [28] was reactivated in both the *nrpe1-L1* and *nrpd1-L1* mutants (Fig S2C), indicating that these two mutations introduce defects in the repression functions of NRPD1 and NRPE1. Thus, we identified NRPE1 and NRPD1 as new components that are involved in repressing the aberrant transcription of *FLC*^*TE*^. This finding, together with previous result [25], suggests that the RdDM pathway can primarily repress intronic TE-mediated aberrant *FLC*^*TE*^ transcription.

The transcription of whole *FLC* locus was modulated by different mechanisms, including FRI-activating and FCA-repressing pathways [29]. Northern blot analysis showed that *FLC*^*TE*^ transcripts were not present in either *Ler, fca* or functional FRI plants (Fig S2D), and neither Mutator transposon-generated siRNA (siRNA^TE^) nor DNA methylation levels were altered in these plants (Fig S3A and S7C), indicating that transcription process of *FLC*^*TE*^ is not modulated by the FRI or autonomous pathway. Both *FLC*^*TE*^ and *FLC* transcripts were higher in *FRI nrpe1-L1* and *FRI ago4* than that in *nrpe1-L1* and *ago4*, respectively (Fig S2D), similar to the comparison of *fca ago4* and *ago4* (Fig S2D). Interestingly, while *FLC*^*TE*^ transcripts increased, the levels of normal *FLC* decreased dramatically in both *FRI* and *fca* backgrounds that carried the RdDM mutants *nrpe1* and *ago4* compared to that in *FRI* and *fca* (Fig S2D). These results suggested that functional *FLC* production was down-regulated with the increasing of intronic *FLC*^*TE*^ in the absence of Pol V- and AGO4-mediated suppression. Consistent with the functional *FLC* expression levels, the flowering time of *fca ago4* plants was earlier than the *fca* plants, and similarly, both *nrpe1-L1 FRI* and *ago4 FRI* dramatically promoted flowering (Fig S2F). While under short-day conditions, *ago4* and *nrpe1-L1* mutants flower earlier than wild type *Ler*, and all these flowering time data demonstrate the aberrant transcription of *FLC*^*TE*^ has important function on plant development.

### Identify the cryptic transcription initiation sites of chimeric *FLC*^*TE*^

While performing northern blot to quantitatively measure the transcript levels of normal *FLC* and aberrant *FLC*^*TE*^, we noticed that when hybridized with Probe B, *FLC*^*TE*^ transcript appeared on top of the blot, with the similar level among *nrpd1-L1, nrpe1-L1* and *ago4* mutants (Fig 1A and S2B). However, surprisingly, when using Probe A, which only contains *FLC* exon 1, the *FLC*^*TE*^ transcript cannot be detected (Fig 1A). As the band corresponding to the normal *FLC* transcript can be detected and present the similar pattern with both Probes A and B in different genetic background (Fig 1A), this data pointed to a possibility that *FLC*^*TE*^ transcript does not contain *FLC* exon 1, which cannot be explained by the previous data and hypothesis [25]. We further assume that, most possibly, it may choose alternative transcription initiation site(s) rather than use the same initiation site as the normal *FLC* transcript.

**Figure 1.**
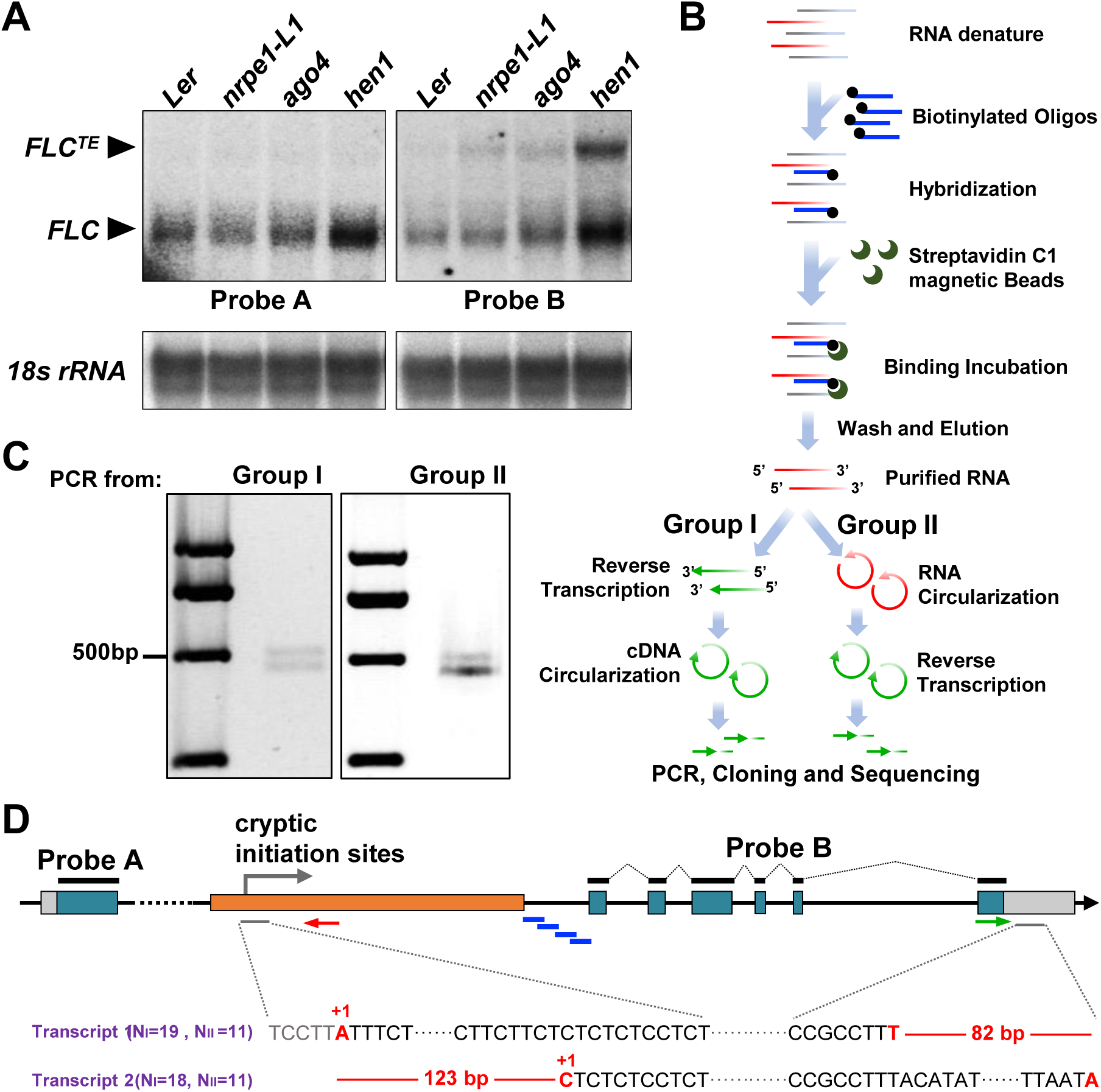
*NRPE1* represses cryptic transcription initiation of *Ler FLC*. (**A**) Aberrant *FLC*^*TE*^ transcripts can be detected in *nrpe1-L1, ago4* and *hen1* mutants using Probe B, but not Probe A, by northern blot analysis. Probe A and B for hybridization regions are shown in Figure 1D. Northern blot of 18S rRNA was used as control. (**B**) Strategies for capturing *FLC*^*TE*^ transcripts and TSS analysis. Four biotinylated single strand DNA probes (light blue lines marked in Figure 1D) specifically mapped on TE-exon2 junction region are used for *FLC*^*TE*^ transcript enrichment. After elution and purification, enriched RNA is respectively subjected to two groups: I) reverse transcription followed by cDNA circularization; II) first RNA circularization and then reverse transcription. Both cDNA templates are used for PCR with a pair of primers shown in Figure 1D (red and green arrows). PCR products are cloned and sequenced. (**C**) Agarose gels results show the same types of PCR products based on cDNA templates from group I and II in Figure 1B. (**D**) Sanger sequencing results of two PCR bands around 500 bp in Figure 2C show that two cryptic initiation sites exist within TE region in *nrpe1-L1* mutant. Five nucleotides before the initiation site A are marked in gray color. N_I_ and N_II_ means the number of sequenced clones from group I and II respectively in Figure 1B.

To test this hypothesis, we had tried hardly with regular 5’ and 3’ end mapping by different regular RACE (Rapid amplification of cDNA ends) methods but none of them worked (data not shown). This might be due to the sequence of the *FLC* intronic TE has multiple copies in the genome and it is hard to detect this low amount of RNA transcripts. Alternatively, through subcellular fraction results, we found that *FLC*^*TE*^ transcripts were highly accumulated in nuclei but not in cytoplasmic fraction (Fig S4A), and the transcripts were dramatically enriched in the on-chromatin fraction (Fig S4B), indicating *FLC*^*TE*^ transcripts are associated with chromatin, and might carry distinct features to the regular RNA molecular forms (which contain caps in 5’ end and polyadenlation in 3’ end). Therefore, we started to enrich *FLC*^*TE*^ transcript with synthesized 5’ biotin labeled DNA probes that can specifically hybridize with *FLC*^*TE*^ exon 2 junction regions (Illustrated in Fig 1D). In order to avoid possibility that the RNA 5’ end structure could affect RNA ligation effeciency, we splited the captured RNAs into two parallel groups (Fig 1B). In Group I, RNA was converted into cDNA followed by circularization; and in Group II, RNA was firstly circularized and then reverse transcribed (Fig 1B). After cloning and sequencing of the DNA products (Fig 1B and 1C), we found that there are two cryptic transcription initiation sites (Transcript 1 and 2, Fig 1D), and both of them localized inner the 5’ end of TE (Fig 1D and Fig S4C). Transcript 1 is initiated from the nucleotide A (data of 19 clones from Group I and 11 clones from Group II), and the initiation core sequence ‘TTA**+1**TTT’ belongs to the previous reported typical and conserved TSS motif in Arabidopsis [30]. Transcript 2 is initiated from the nucleotide C (18 clones from Group I and 11 clones from Group II), and the initiation core sequence ‘CTC**+1**TCT’ is atypical (Fig 1D and Fig S4C). Additionally, these two initiation sites (A to C) has 123 bp in distance, where their termination sites on the 3’ UTR of normal *FLC* vary 82 bp in distance (Fig 1D and Fig S4C). Interestingly, the sequencing data from either circularized cDNA or RNA did not observed the poly A sequence, suggesting the chimeric *FLC*^*TE*^ transcripts are not polyadenylated, consistant with its chromatin association feature (Fig S4A and S4B). In addition to the initiation and termination variation, we also confirmed two splicing sites around *FLC*^*TE*^ exon 2 in *nrpd1-L1* and *nrpe1-L1* mutants from RT-PCR results (Fig S2C and S4B), indicating that splicing events may happen during *FLC*^*TE*^ RNA processing. Taken together, we have identified the cryptic transcription initiation sites of chimeric *FLC*^*TE*^, which is necessary for functional study of *FLC*^*TE*^ biogenesis and regulatory mechanism.

### Intronic heterochromatin and DNA methylation suppress the *FLC*^*TE*^ cryptic initiation

To understand the mechanism how to initiate the cryptic *FLC*^*TE*^ transcription, we first analyzed DNA methylation levels on *FLC* Mutator region, and they were hardly detected in two mutants *nrpe1-L1* and *nrpd1-L1* (Fig 2B and S7A). Then we checked histone repressive mark H3K9me2, for most of RdDM-repressed genome loci are associated with it [23,24]. Chromatin immunoprecipitation (ChIP) results show that H3K9me2 levels on the *FLC* Mutator region were dramatically reduced in *nrpe1-L1, nrpd1-L1* and *ago4* compared to *Ler* (Fig 2C), indicating that the intronic heterochromatin formation on the TE region depends on RdDM pathway. Transcription initiation process is tightly determined by chromatin states which control the entry of initiation form of RNA polymerase II (Pol II). Coupled with chromatin state, the phosphorylation on Serine 5 of RNA Pol II CTD (carboxyl terminal domain) repeats (Pol II Ser5p) are essential to regulate transcription initiation [31-33]. The level of Pol II Ser5p on TE and its neighbor regions was significantly higher in *nrpe1-L1* than in *Ler* (Fig 2D), suggesting that the defects in intronic heterochromatin formation provide opportunity for Pol II to entry and recognize the cryptic initiation sites. Our results also show that H3K4me3, which normally represented as the transcription initiation mark [34], was exclusively enriched on TE region close to cryptic initiation sites in *nrpe1-L1* (Fig 2E). The unprocessed *FLC* transcripts were highly enriched in *nrpe1-L1* and *ago4*, especially in the region downstream of *FLC*^*TE*^ initiation sites (Fig S5A). We also found the elongated form of Pol II (Pol II Ser2p) was significantly enriched in *nrpe1-L1* than in *Ler* (Fig S5B), nuclear run-on assay showed the transcription elongation rate was increased in *nrpe1-L1* (Fig 2G), and the transcription elongation mark H3K36me3 deposited in higher level in *nrpe1-L1* than in *Ler* (Fig S5C). Together with results that total Pol II occupancy was also significantly enriched on *nrpe1-L1 FLC* locus (Fig S5D), when depleting intronic heterochromatin, the cryptic transcription initiation sites are preferably recognized by Pol II accompany with transcription initiation and fast elongation (Model Fig 2H). While observing the RdDM-dependent heterochromatin establishment, we hypothesized that the *FLC* intronic heterochromatin could be the direct target of RdDM such as NRPE1. Previous genome-wide ChIP-seq clearly showed that NRPE1 is associated with multiple heterochromatin loci [35,36], including the homologous region of *FLC*^*TE*^ Mutator in Col-0 (Fig S8). Indeed, our ChIP results also showed the clear enrichments of NRPE1 around the Mutator region in *FLC* (Fig 2F and S5E).

**Figure 2.**
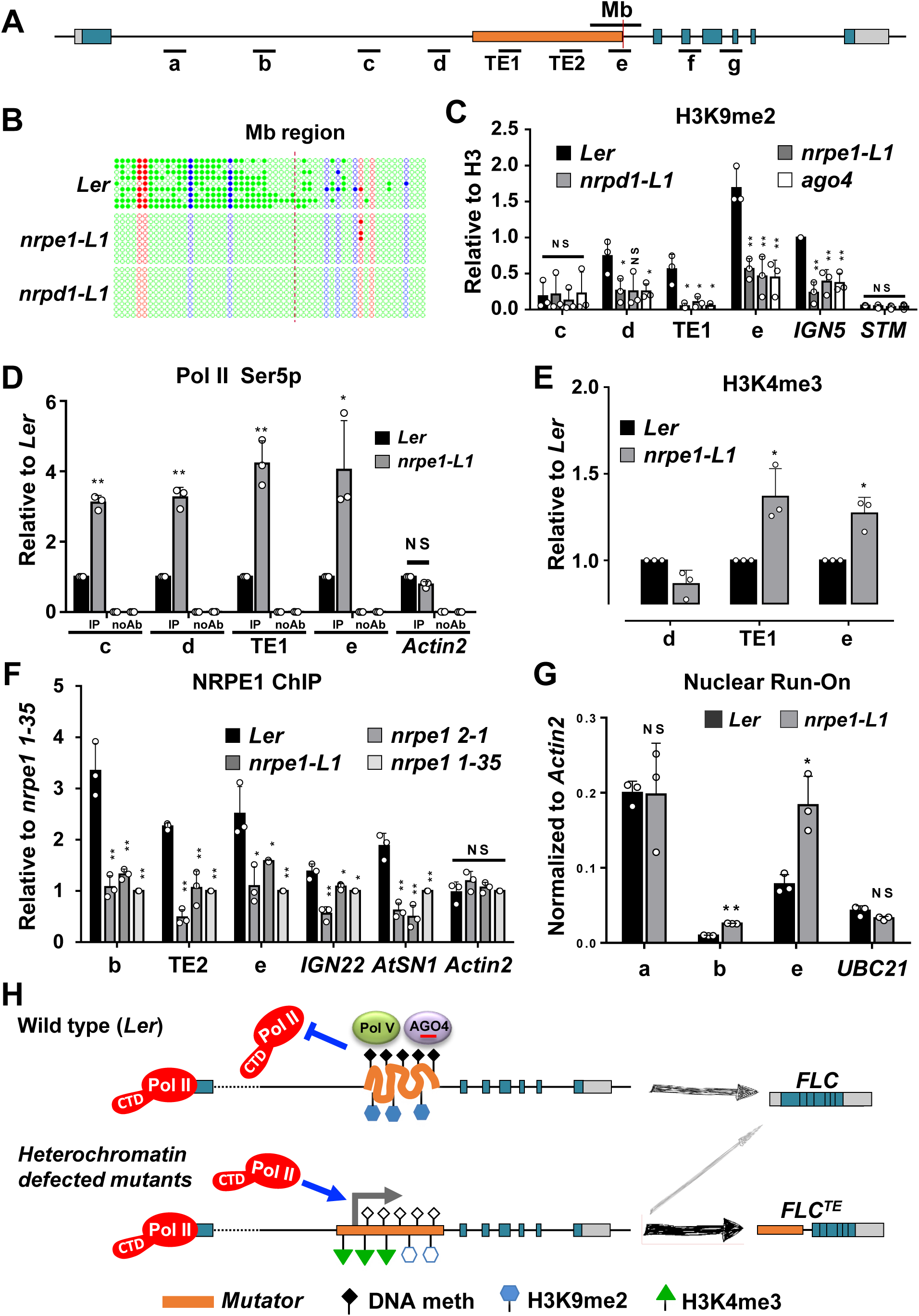
Repression of *FLC* cryptic initiation requires the NRPE1-dependent intronic heterochromatin formation. (**A**) Schematic representation of *FLC* locus marked by qPCR primers and bisulfite sequencing region. (**B**) DNA methylation levels on the sense strand of *FLC* (region Mb) was analyzed by bisulfite sequencing. Red dashed line marks the right border of the TE region. Quantitive accumulation of CG, CGH and CHH methylation levels (%) on region Mb are shown in Figure S6A. (**C**) H3K9me2 enrichement on TE reguion is reduced in *nrpe1-L1, nrpd1-L1* and *ago4*. ChIP values were normalized to H3 signals. *ING5* and *STM* were used as positive and negative controls, respectively. (**D**) ChIP-qPCR of Pol II Ser5p shows increasing the Pol II initiation in *nrpe1-L1* mutant compared to *Ler. Actin2* was used as negative control. (**E**) Transcription initiation marker H3K4me3 on TE region is increased in *nrpe1-L1* mutant. (**F**) ChIP-qPCR analysis of NRPE1 protein occupancy on *FLC* using anti-NRPE1 antibody. *IGN22* and *AtSN1* were used as positive controls, while *Actin2* was used as a negative control. Mutants of *nrpe1 1-35* and *2-1* are two lines from independent crossing of *Ler* with *nrpe1* (Col-0 background). (**G**) Nuclear run-on analysis to detect elongation rates of Pol II in *Ler* and *nrpe1-L1.* RNA levels were normalized to *Actin2. UBC21* was used as a negative control. (**H**) A working model describes how the heterochromatin prevents intronic transposon-mediated cryptic transcription initiation on *FLC* locus. In WT *Ler*, Pol V and AGO4 associate with local heterochromatin restricted to the *FLC* Mutator transposon element, where DNA methylation, H3K9me2 and siRNA are highly enriched. RdDM-dependent intronic heterochromatin inhibits Pol II entry into the potential transcription initiation sites within TE region, resulting in only transcribing normal *FLC*. In heterochromatin defected mutants such as *nrpe1* or *ago4*, the depletion of DNA methylation and H3K9me2 makes TE chromatin more accessible, coupled with increased H3K4me3 and H3K36me3, resulting in recognizing cryptic transcription initiation by Pol II. Under this condition, the on-chromatin *FLC*^*TE*^ transcript is synthesized to repress the normal *FLC*. Curved and linear TEs (shown in orange) represent condensed heterochromatin and opening euchromatin, respectively.

We then examined the *FLC*^*TE*^ transcription initiation role of other canonical RdDM pathway components that involved in siRNA biogenesis, scaffold RNA transcription, *de novo* DNA methylation, chromatin remodeling and heterochromatin formation [23,24]. *FLC*^*TE*^ transcripts were significantly elevated in *nrpd1, rdr2* and *dcl3* (Fig S6C), and consequently, siRNA^TE^was undetectable in *nrpe1-L1, ago4, nrpd1, rdr2* and *dcl3* (Fig S6D). *FLC*^*TE*^ transcripts in *ktf1* mutant strongly accumulated to the comparable levels found in *nrpe1-L1* and *ago4* (Fig S6C). However, siRNA^TE^levels were not altered in the *ktf1* mutant compared to other RdDM mutants (Fig S6D), which was consistant with the previous results that *ktf1* mutant has no effect on the siRNAs accumulation from RdDM targets [37]. We also found the disconnection between siRNA^TE^ and *FLC*^*TE*^ production in different tissues (Fig S3B and S3C). These data suggested that the suppression of cryptic initiation does not rely on local siRNA^TE^ biogenesis.

DNA methylation levels were dramatically reduced in *nrpe1-L1, ago4, nrpd1, rdr2, dcl3, ktf1* and *drd1* mutants (Fig S7B). DRM2 is the main DNA methyltransferase in the RdDM pathway that catalyzes *de novo* CHH methylation [23,24], although it functions redundantly with DRM1 [38-40]. Indeed, our results showed that CHH DNA methylation of *FLC* Mutator was strongly reduced in the *drm2* mutant (Fig S6A), but not fully reduced to *nrpe1-L1, ago4* or *drm1drm2* levels (Fig S6A), and the dramatic DNA methylation reduction happened in the *drm1drm2* double mutant compared to *nrpe1-L1* and *ago4* (Fig S6A). We found no strong increase in *FLC*^*TE*^ transcripts in either the *drm1* or *drm2* single mutants compared to *Ler* plants (Fig S6B). Interestingly, a clear increase in *FLC*^*TE*^ transcripts abundance in the *drm1drm2* double mutant to the levels comparable to, or slight higher than, those in *nrpe1-L1* and *ago4* (Fig S6B). The results demonstrate that DNA methylation (especially CHH) that was catalyzed by both DRM1 and DRM2 is essential for *FLC*^*TE*^ transcription initiation.

Taken together, RdDM pathway is essential to repress *FLC*^*TE*^ transcription initiation at the cryptic sites in the intronic Mutator transposon. The accessible chromatin that is essential for the new transcription initiation can be switched to intronic inaccessible heterochromatin through the activity of RdDM pathway (Fig 2H). Therefore, we emphasize the notion that the chromatin state determines the recognition of cryptic initiation sites in introns which could extend from *Ler FLC* locus to other genomic loci.

### RdDM-suppressed cryptic initiation at intronic heterochromatin is a common mechanism in the Arabidopsis genome

In order to inverstigate what we had discovered on *FLC* locus is commonly happened in other genomic loci, we started to profile the Pol II initiation state in genome-wide level with the established pNET-seq (plant native elongating transcript sequencing [41]) method by using RNA Pol II Ser5p antibody in wild type Col-0 and *nrpe1* mutant (Fig S9). To precisely understand the mechanism of RdDM mediated suppression of cryptic initiation, we also included the previous published H3K4me2 and -me3 ChIP-seq, whole genome bisulfite-seq, and NRPE1 ChIP-seq and RIP-seq data for genome wide analysis [42-44].

In total, there are 523 upregulated Ser5p peaks (US5Ps) identified in *nrpe1* mutant compared to Col-0 based on our pNET-seq data, and most of the US5Ps are enriched in intragenic regions (Fig 3A and 3B). Interstingly, these US5Ps significantly presented in association with higher H3K9me2 levels than the downregulated Ser5p peaks (DS5Ps) (Fig 3C), indicating that the new RNA transcripts are prior to be reinitiated from the repressive heterochromatin regions in the absence of NRPE1. Consistant with DNA methylation patterns in *nrpe1* mutant, we detected significant loss of DNA methylation levels in all sequence contexts (CG, CHG and CHH) on US5Ps but not on DS5Ps (Fig 3D and 3E), which implies that these DS5Ps could not be directly targeted by RdDM pathway. Moreover, only US5Ps have elevated active transcription initiation marks H3K4me2/3 (Fig 3D and 3E). These results indicate that the reactivation regions in the *nrpe1* mutant are regulated by loss of DNA methylation and deposition of H3K4me2/3 marks, which is consistant with the case of *Ler FLC* that we had observed above (Fig 2B, 2D and 2E). Then we checked whether these US5Ps were directly targeted by Pol V or not [35], the results show that more than 30% of US5Ps are directly associated with NRPE1 (158 in 523 peaks, Fig 4A). Consistant with the key function of NRPE1 in RdDM pathway, DNA methylation in all contexts on these Pol V associated US5Ps are reduced (Fig 4B). Moreover, H3K4me2/3 modifications are increased significantly (Fig 4B), which suggests that reactivation of Pol V associated US5Ps is induced by loss-of-RdDM-dependent H3K4me2/3 deposition. Although we oberserved conspicuous decrease of DNA methylation levels on those non-Pol V associated US5Ps (Fig S10A), no significant increase of H3K4me2/3 levels can be detected on non-Pol V associated US5Ps (Fig S10A), which indicates that reactivation of non-Pol V associated US5Ps in *nrpe1* is not associated to H3K4me2/3 modifications. In addition, among 492 Pol V associated introns, 247 upregulated introns with Ser5 signals are marked in *nrpe1* mutant (Fig 4C), and as expected, they obviously showed the loss of DNA methylation and increase of H3K4me2/3 modificaitons (Fig 4D). In contrast, other introns do not have these reactivation featrures (Fig S10B). Among these upregulated introns, 30 introns are observed to contain annotated TEs and these intronic TEs are distributed in various subfamilies (Table S2). RT-PCR results from two selected genes (AT5G52070 and AT3G33528), which have the increased Ser5p signal and loss of CHH methylation in introns, also confirm the new intronic initiations that truly exist in *nrpe1* mutant (Fig 5). Together, these results suggested the intronic heterochromatin-repressed cryptic initiation is a common phenomenon in Arabidopsis.

**Figure 3.**
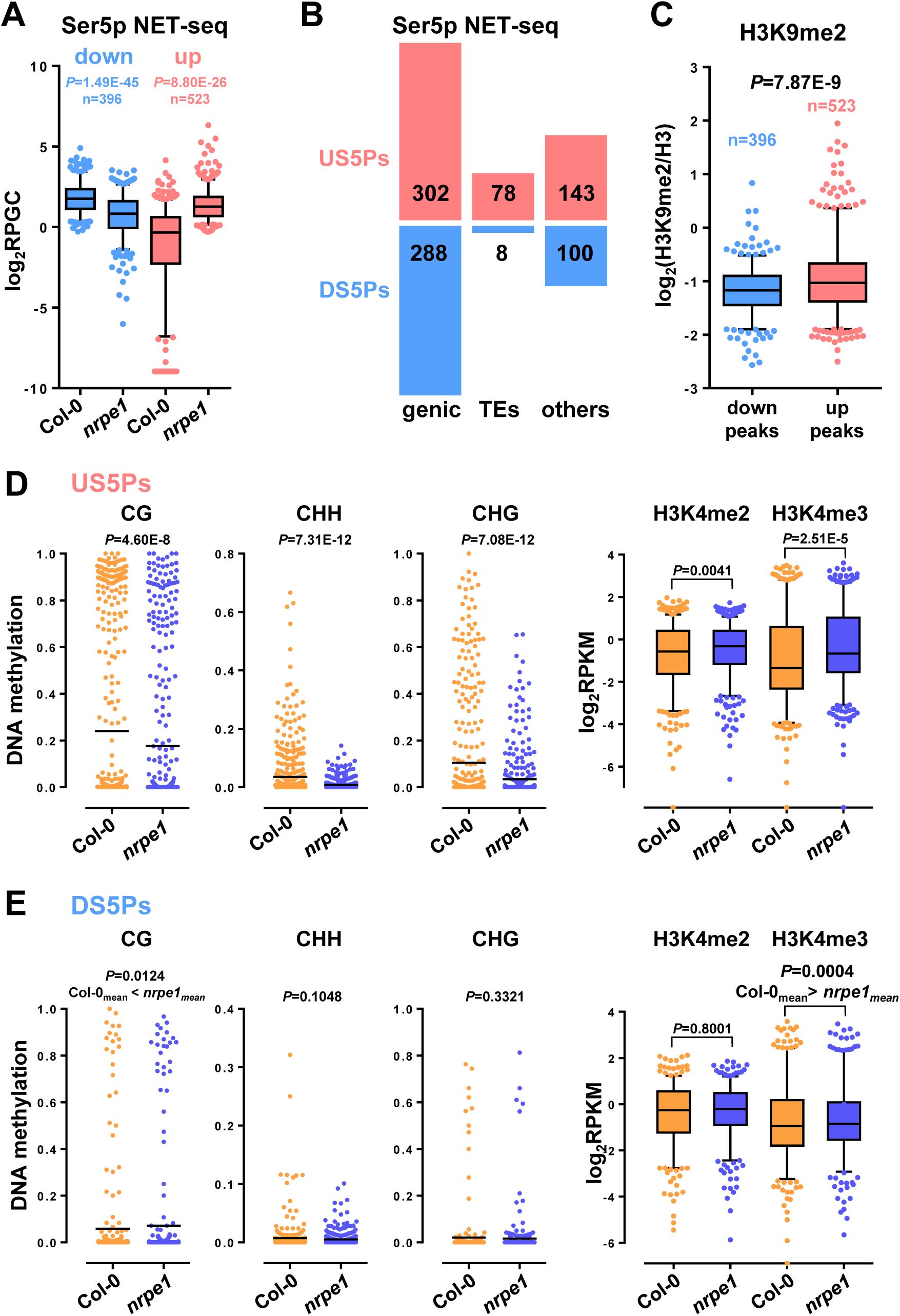
Cryptical transcription initiations are widespread in *nrpe1* mutant. (**A**) Box plots show Pol II Ser5p pNET-seq values of differentially expressed pNET-seq peaks in *nrpe1* versus Col-0. Values are the mean of RPGC counts from two replicates. (**B**) The bar plots show the number of differentially expressed pNET-seq peaks in *nrpe1* versus Col-0 in classified genomic regions. (**C**) Box plots show H3K9me2 levels of differentially expressed pNET-seq peaks in *nrpe1* versus Col-0. Values are the mean of RPKM counts from two replicates then normalized to mean of H3 RPKM counts. (**D and E**) DNA methylation levels and H3K4me2/3 levels of upregulated or downregulated Ser5p pNET-seq peaks in *nrpe1* versus Col-0.

**Figure 4.**
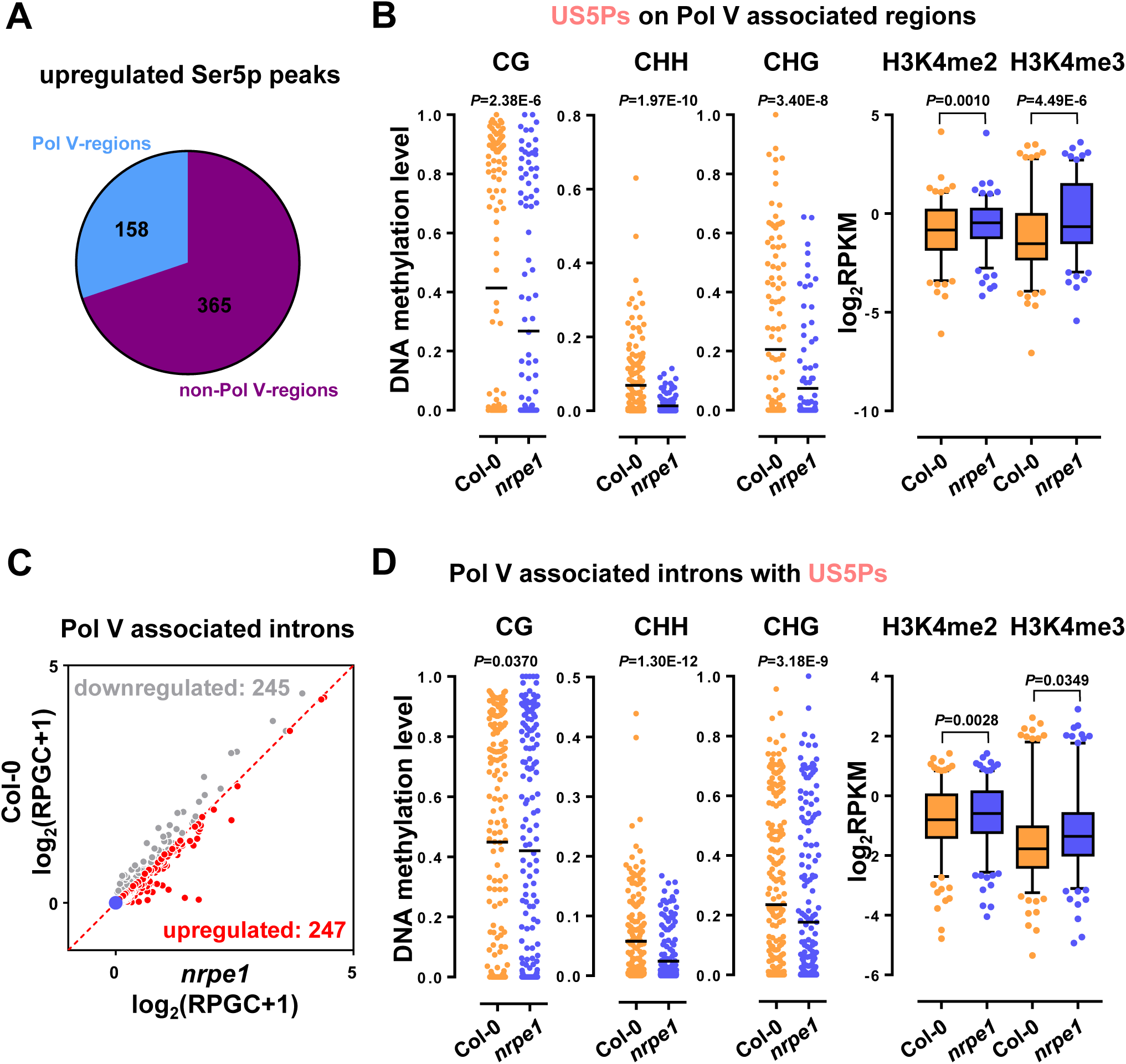
Pol V associated intronic initiations are triggered by loss of DNA methylation and deposition of H3K4me2/3 modifications. (**A**) The number of upregulated Ser5 pNET-seq peaks in *nrpe1* versus Col-0 on Pol V or non-Pol V associated regions. (**B**) DNA methylation levels and H3K4me2/3 levels of on Pol V associated regions. (**C**) Scatterplots of different regulated Pol V associated-introns from pNET-seq data in *nrpe1* versus Col-0. Values are the mean of log2(RPGC counts +1) from two replicates. (**D**) DNA methylation levels and H3K4me2/3 levels of upregulated Pol V associated-introns according to Ser5p pNET-seq data.

**Figure 5.**
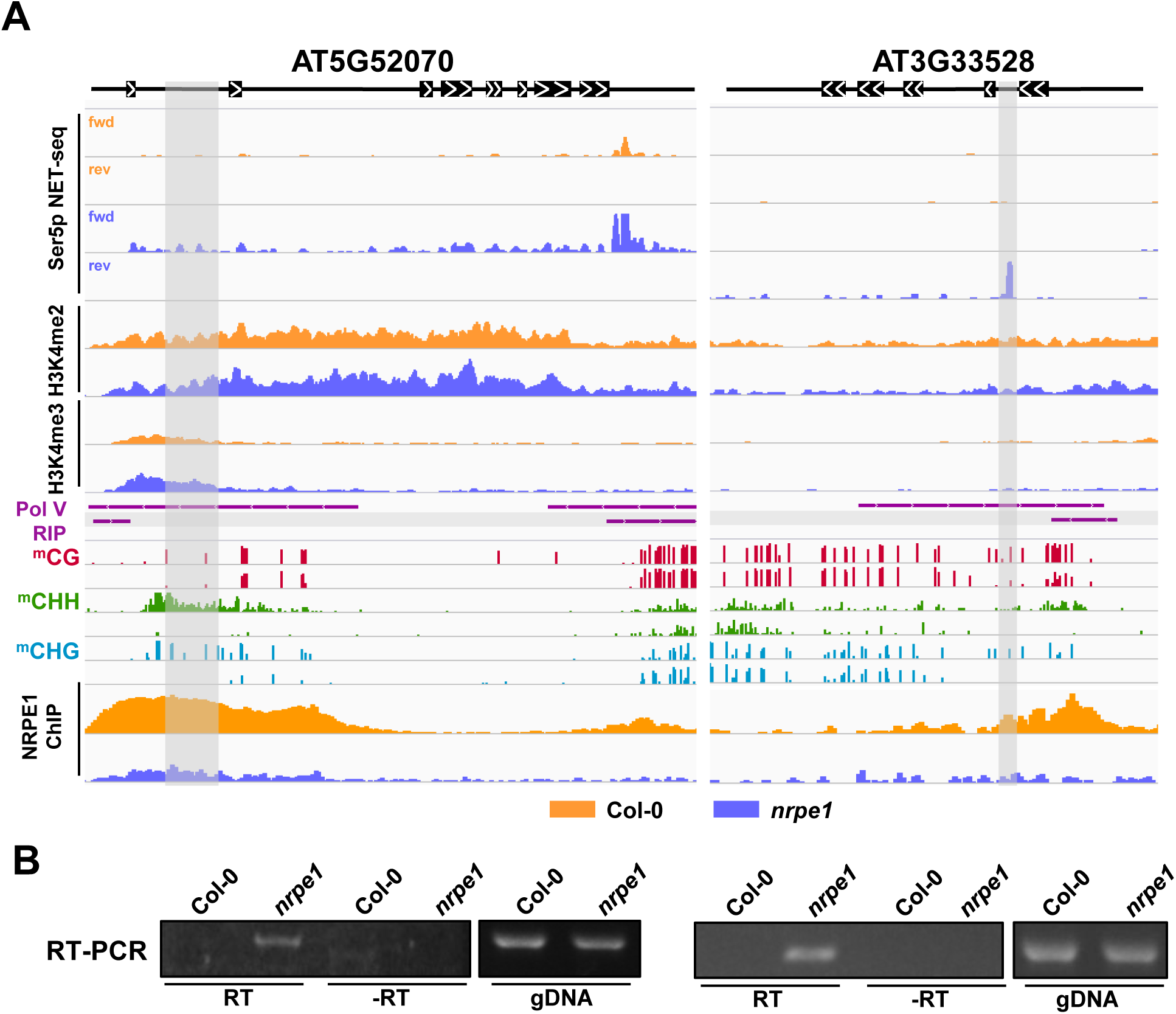
Two examples of Pol V-repressed intronic initiations. (**A**) Snapshot of indicated genes with Ser5 pNET-seq, H3H4me2/3, Pol V RIP-seq, whole-genome bisulfite sequencing and NRPE1 ChIP-seq data. (**B**) RT-PCR assays to comfirm innitations on indicated genes. Amplification regions are indicated with gray boxes in Figure 5A.

## DISCUSSION

TEs are abundantly present in various locations in the plant genome [2]. Most studies focus on non-genic TEs and their regulatory mechanisms, while less is known about the function of genic TEs which comprise a large proportion in the genomes. Using *FLC* in *Ler* ecotype that carries an intragenic TE as a model system, combining with forward genetic screening, we revealed a novel regulatory mechanism that RdDM-mediated heterochromatin formation on *FLC* intronic TE suppresses cryptic transcription initiation locally. The intronic heterochromatin maintained by DNA methylation and H3K9me2 prevents initiation sites recognition by RNA Pol II machinery. Our work uncovered a new function of RdDM pathway on intronic TE is important for repressing *FLC*^*TE*^ transcription initiation (Fig 2H).

Removing DNA methylation and heterochromatic marks in different RdDM mutant backgrounds increased the occupancies of initiating form of Pol II Ser5p and H3K4me3 nearby the cryptic initiation sites (Fig 2D and 2E), suggesting that defects in heterochromatin promotes RNA Pol II entry and transcription initiation. Since transcription process includes initiation, elongation, termination and recycling steps, transcription elongation is also coupled to determine initiation [45,46]. We observed the increasing enrichment of elongating form of Pol II Ser2p as well as elongation coupled epigenetic modification mark H3K36me3 on *FLC* (Fig S5B and S5C). Related to our observation, histone methyltransferase SETD2 recruited by elongating RNA Pol II to catalyze H3K36me3 which is essential for gene-body DNA methylation and suppression of cryptic initiation [47].

Multiple pathways regulate *FLC*, which encodes a central repressor of the floral transition [29,48-50]. Even though FRI and FCA do not regulate *FLC*^*TE*^ transcription genetically (Fig S2D and S2F), *FLC*^*TE*^ transcript levels dramatically increased in RdDM mutants with both *FRI* and *fca* backgrounds (Fig S2D). We also observed early flowering in *nrpe1-L1* and *ago4* mutants in both *FRI* and *fca* backgrounds, or under short-day conditions (Fig S2F and S2G). In line with the phenotype, expression levels of *FLC* are reduced in RdDM mutants combined with *FRI* and *fca* backgrounds (Fig S2D). These results suggested that the chimeric *FLC*^*TE*^ transcripts accumulation could negatively regulates functional *FLC* production, most possibally through competing different initiation sites in the same locus (Fig 2H).

The intronic heterochromatin in *IBM1* gene can modulate its mRNA termination and processing (polyadenylation), which was altered by disturbed interactions between RNA-binding protein IBM2 and heterochromatin maintenance, but without DNA methylation alteration [17,18]. In contrast, our results showed that, in the absence of RdDM and DNA methylation, the NRPE1-associated intronic heterochromatin regions accelerate the transcription initiation with depositing the initiation marks H3K4me2 and -me3 (Fig 4B and 4D), thus results in initiating the new transcription. Together with published results that H3K4 demethylases LDL1 and LDL2 work together to maintain RdDM patterns [42] and gene-body H3K4me1/H3K9me2 dynamics are precisely modulated by two opposing activities of histone demethylases LDL2 and IBM1 [51], we conclude that H3K4me and heterochromatic modifications (DNA methylation and H3K9me2) have antagonistic roles for regulating transcription initiation on specific intronic TE regions, probably through H3K4me3 methyltransferases activity, such as ATX1, ATX2, ATXR3, and ATXR7 [52]. However, further study is still required to identify which is the major methyltransferase that is responsible for depositing H3K4 methylation for promopting intiation in the case of RdDM depleting.

R-loop, which is comprised of a DNA-RNA hybrid and the displaced single-stranded DNA, was reported to modulate *FLC* expression through modulating the activity of long non-coding RNA *COOLAIR* [53]. In this study, we also noticed R-loop features on Ser5p upregulated and Pol V associated intronic peaks, because the purine Guanine and Adenine rich motif was found on the top of these Ser5p upregulated intronic peaks (Fig S11A). Our previous study had already revealed a similar “GARGAAGx” (where R = purine G or A) motif on the three stranded chromatin structure R-loops in Arabidopsis [54]. R-loop prediction in mammalian genome also pointed out the purine-rich sequences, such as GA motif, are prone to forming DNA:RNA hybrids [55]. Interestingly, our previous observations that R-loops in Arabidopsis genome are highly associated with heterochromatin marks H3K9me2 and Pol IV loci, but not Pol V loci [54]. In addition, our recent work also showed that a histone-like protein complex specially reads genic R-loops, and depletion of this complex does not alter genome-wide R-loop levels but sensitive to stress stimulation (Yuan *et al.*, ***Science Advances***, accepted). Thus, future studies may also be required to understand the intragenic R-loops structure in platforming the transcription regulation under various conditions.

Together, the mechanistic dissection on the intronic heterochromatin-containing genes refreshes our understanding of genic TEs in regulating their host genes, and pointed out the alternative possibilities that intronic TEs formed heterochromatin could potentially be used as cryptic initiation sites while in absence of the suppression components (this study) or during stimulations, and many chromatin states such as transcription activation marks and R-loops are tightly involved.

## Materials and Methods

### RT-qPCR-based forward genetic screening and fine mapping of *nrpd1-L1* and *nrpe1-L1*

The mutant *nrpe1-L1* was isolated from an EMS-mutagenized population on the *Ler* background. Briefly, after RNA extraction from all seedlings from an EMS-mutagenized population, we performed reverse transcription using gene-specific primers (5’-CTTTTGACTGATGATCCAAGGCTTTAA-3’), followed by quantitative PCR (qPCR) using *FLC*^*TE*^ detection primers (5’-CTCTCCTCTCCGCCTTTCTC -3’ and 5’-TGTGGTGAGCCTGTAACTATGAA-3’). Two candidates (*mutant a* and *b*, Fig S1A and B) with more than 20-times the signal strength of wild type *Ler* were crossed with Col-0, and F2 plants were used for mapping. More information for mutants mapping can be found in Fig S1.

### Plant materials and growth conditions

The mutants *nrpe1* (SALK_092219), *nrpd1a* (SALK_128428), *rdr2* (SAIL_1277_H08), *dcl3* (SALK_005512), *drd1* [56], *ktf1* [37], *rdr6* and *drm1drm2* [39] were crossed with *nrpe1-L1* or *ago4-1* to generate homozygous mutants with a diploid *Ler-FLC* locus. *FRI Ler* or *fca* in the *Ler* background were crossed with *nrpe1-L1* or *ago4-1* to generate homozygous double mutants. All primers used for genotyping were adapted from the original paper (detailed information is in Table S1) and are summarized in Table S1. All plants were grown on 1/2 MS medium plates for 10-14 days under long-day conditions (16 h light at 22°C, 8 h darkness at 20°C) after 2 days of stratification at 4°C. For the short-day flowering time measurement, growing conditions are 8 h light at 22°C, 16 h darkness at 20°C.

### *FLC*^*TE*^ transcripts enrichment and TSS analysis

This method was modified from ChIRP [57]. Total RNA (∼1 mg) extracted from Trizol reagent was dissolved in 1 mL Hybridization Buffer (50 mM Tris-HCl pH7.0, 750 mM NaCl,1% SDS, 1 mM EDTA, 15% formamide, 0.1 mM PMSF and 0.1 U/µl RNase inhibitor) and then was denatured for 15 min at 55°C, then immediately chilled on ice for at least 1 min. 10 μl biotinylated DNA probe pool (5’Biotin-aatttccactttttcaaaaaagttttaacttgttcttagctttgtaagcaatatttaat -3’, 5’Biotin-attcgtaacatttcagaaaaacaataatcttaatattcatgtttaagcagtagcacatc -3’, 5’Biotin-tgaaaaggccactggaaactatgaaacattgagagaacaccttaccttttacatatata -3’, 5’Biotin-cttgaccaggctggagagatgacaaaaaaaataaaaatatgttaggatcaaaactacta -3’, 10mM for each probe) was also heat-denatured (15 min at 55°C) and chilled on ice immediately. Add denatured DNA probe pool to denatured RNA and transfer to a 37°C hybridization oven and incubate for 4 hours with gentle rotation. Wash 20 μl of Streptavidin C1 (Invitrogen, 65001) before use: wash twice in 500 μl buffer I (0.1 M NaOH, 0.05 M NaCl) for 2min and wash once in 500 μl buffer II (0.1 M NaCl), final once wash in 500 μl Hybridization Buffer. Pre-washed beads were resuspended in 20 μl Hybridization Buffer and then added into RNA-probe mixture. After incubation at 37°C for 30min with rotation, RNA-beads complex was washed twice with 37°C pre-heated Wash Buffer A (2XSSC, 0.5% SDS, 1mM DTT and 1mM PMSF) and was washed twice with 37°C pre-heated Wash Buffer B (0.1XSSC, 0.5% SDS, 1mM DTT and 1mM PMSF). For target RNA elution, resuspended the beads in 200μl Elution Buffer (100 mM NaCl, 50 mM Tris-HCl pH7.0, 1 mM EDTA, 1%SDS) and boil at 95°C for 15min, then followed by RNA extraction with Trizol reagent.

Extracted RNA from above steps was divided into two parts. One was reverse transcripted with Reverse Transcriptase SuperScript III (Invitrogen) using random primers and then treated with CircLigase (Epicentre). Circular cDNA was used as template for PCR with a pair of primers shown in Fig 1D (red primer: 5’-GTAGATATGTGGTGAGCCTGTAACTATGAAGAAC-3’, green primer: 5’-GAATAATCATCATGTGGGAGCAGAAGCTGAGATG-3’). Another part of enriched RNA was first treated with RppH (NEB, M0356S) and then followed ssRNA Ligase (NEB, M0204) treatment. Circular RNA was reverse transcripted with Reverse Transcriptase SuperScript III (Invitrogen) using random primers and produced cDNA was used as template for PCR with same primers. Both PCR products from above two methods were purified for cloning with pEASY-Blunt Zero Cloning Kit (Transgen Biotech, CB501) and finally performed Sanger sequencing.

### Gene expression analysis

RNA extraction, splicing-specific RT-PCR (ssRT-PCR), and unprocessed transcript level measurements were performed as described below. For ssRT-PCR and unprocessed *FLC* transcript level measurements, total RNA was extracted from the tissue using Trizol reagent followed by DNase I (Ambion, M1903) treatment. Reverse transcription was performed with Reverse Transcriptase SuperScript III (Invitrogen) using gene-specific primers (Table S1). *FLC*^*TE*^ and *FLC* were amplified using the primers shown in Table S1.

### DNA methylation analysis

Two methods, Chop-PCR and bisulfite-sequence, were used to measure DNA methylation levels in the mutants. Genomic DNA was extracted using the CTAB method. For Chop-PCR, 1.5 μg genomic DNA was digested with methylation-sensitive (Alu I, Aci I, Hha I) or methylation-specific restriction enzymes (McrBC), followed by PCR. Non-digested DNA was amplified as a control. For bisulfite sequencing of the *FLC*-TE region, 2 μg genomic DNA was subjected to bisulfite conversion using an EZ DNA Methylation-Gold Kit as described by the manufacturer (D5005). The converted DNA was amplified with primers for the Mutator TE region shown in Table S1. The PCR products were purified using a HiPure Gel Pure DNA Mini Kit (Magen, D2111) and cloned into the pGEM-T Easy Vector (Promega, A1360). For each sample, appropriately 10-15 individual clones were sequenced. Data were analyzed using the online software program Kismeth (http://katahdin.mssm.edu/kismeth/revpage.pl).

### Nuclear run-on assay

14-day-old seedlings were ground into a fine powder in liquid N2 and resuspended in 5 ml pre-cooled nuclease-free Honda buffer (0.44 M sucrose, 1.25% Ficoll, 2.5% Dextran T40, 20 mM HEPES KOH, pH 7.4, 10 mM MgCl2, 0.5% Triton X-100, 5 mM DTT, 1 mM PMSF, 1X Protease Inhibitor). After filtering through two layers of Miracloth and centrifuging at 2000 g for 10 min at 4°C, nuclei were washed 2–3 times with Honda buffer and resuspended in 50 µl nuclei storage buffer (50 mM Tris-HCl, pH 7.8, 10 mM 2-mercaptoethanol, 20% glycerol, 5 mM MgCl2 and 0.44 M sucrose). The run-on assay was performed in a reaction containing 10 µl 10X transcription assay buffer (500 mM pH7.5 Tris-HCl, 50 mM MgCl2, 1.5 M KCl, 1% sarkosyl, 20 U/ml RNase inhibitor [Invitrogen] and 100 mM DTT), 50 µl nuclei in storage buffer, 5 µl NTP mixture (100 mM ATP, 100 mM CTP, 100 mM GTP, 100 mM BrUTP [Sigma, B7166]) and 35 µl H2O. The run-on reaction was performed at 30°C for 30 min. The reaction was stopped by adding 600 µl Trizol reagent (Invitrogen), and RNA was extracted followed by DNase I (Ambion, M1903) treatment. The purified RNA was diluted in 500 µl incubation buffer (20 mM Tris-HCl, pH 7.5, 4 mM MgCl2, 0.2% NP40) and incubated with 2 µl anti-BrdU antibody (Sigma, B8434) at 4°C for 2 h, followed by immunoprecipitation for 1 h with Dynabeads Protein G (Invitrogen). The precipitated RNA was extracted with Trizol reagent and used for RT-qPCR analysis.

### Nuclear-Cytoplasmic Fractionation

The Nuclear-Cytoplasmic Fractionation assay was performed as described [58] with some modifications. Briefly, 14-day-old seedlings (2 g) were ground into fine powder in liquid N2 and resuspended in 10 ml pre-cooled lysis buffer (20 mM pH 7.4 Tris-HCl, 25% Glycerol, 20 mM KCl, 2 mM EDTA, 2.5 mM MgCl2, 250 mM Sucrose, and 5 mM DTT) containing 40 U/ml RNase inhibitor (Invitrogen, AM2696). After filtering through two layers of Miracloth, the flow-through was centrifuged at 1500 g for 10 min. The pellet (containing the nuclear fraction) and the supernatant (containing the cytoplasmic fraction) were transferred to a new tube, centrifuged at 13,000 g for 15 min, and collected. The cytoplasmic RNA was immediately isolated with Trizol reagent. The pellet was washed 4–5 times with pre-cooled NRBT buffer (20 mM pH 7.4 Tris-HCl, 25% Glycerol, 2.5 mM MgCl2, 0.2% Triton X-100, 5 mM DTT and 160 U/ml RNase inhibitor), resuspended in 300 µl pre-cooled Buffer A (250 mM sucrose, 10 mM Tris-HCl, pH 8.0, 10 mM MgCl2, 1% Triton X-100, 5 mM 2-mercaptoethanol, 1X Protease Inhibitor and 350 U/ml RNase inhibitor) and layered on top of 300 µl pre-cooled Buffer B (1.7 M sucrose, 10 mM Tris-HCl pH 8.0, 2 mM MgCl_2_, 0.15% Triton X-100, 5 mM 2-mercaptoethanol, 1X Protease Inhibitor and 350 U/ml RNase inhibitor) in a new tube. After centrifugation at 13,000 g for 45 min at 4°C, the pellet was subjected to nuclear RNA extraction with Trizol reagent. Nuclei fractionation was performed as previously described [59].

### Northern blot analysis

Total RNA was extracted from Arabidopsis seedlings or tissues using Trizol reagent (Invitrogen). Approximately 20 µg of RNA was separated on a 1.5% agarose-formaldehyde denaturing gel and transferred onto nylon membranes (Amersham). Hybridization was performed in SuperHyb™ hybridization buffer (Sigma) for 16 h at 68°C. The membranes were washed twice for 5 min with 2X SSC and 0.1% SDS, washed twice for 15 min with 0.1X SSC and 0.1% SDS and exposed to a phosphorimager screen for 2–3 days. Specific DNA probes against sense *FLC* were labeled with α-^32^P-dCTP in PCR with gene-specific reverse primers. PCR amplifications of DNA probe a and b were performed using the gene-specific primers listed in Table S1.

Northern blot analysis of small RNA was performed according to previous procedures [60]: 20–30 μg small RNA was enriched from total RNA using PEG4000, separated on a denaturing gel and transferred onto nylon membranes (Amersham). Hybridization was performed in hybridization buffer (1% BSA [w/v], 0.5 M Na_2_HPO_4_, 1 mM EDTA, 7% SDS, 15% formamide [v/v]) for 16 h at 42°C. The membranes were washed three times for 10 min per wash with 1X SSC and 0.1% SDS. DNA Probes against the transposon sequence on *FLC* were prepared according to standard northern blot procedures. PCR amplification of the TE region was performed using the gene-specific primers listed in Table S1.

### Chromatin immunoprecipitation (ChIP)

For the NRPE1-Flag and native NRPE1 ChIP, seedlings grown on solid 1/2 MS medium were harvested in PBS buffer with 1% formaldehyde (Sigma). The plants were treated under a vacuum for 10 minutes, and glycine (0.125 M) was added to terminate the formaldehyde crosslinking reaction. The samples were ground to a fine powder in liquid nitrogen, and debris was removed by filtering the samples through Miracloth (Merck). Nuclei were isolated in Honda Buffer comprising 0.44 M sucrose, 1.25% Ficoll, 2.5% Dextran T40, 20 mM HEPES (pH 7.4), 10 mM MgCl_2_, 0.5% Triton X-100, 5 mM DTT and 1X protease inhibitor (Roche). Loose chromatin was removed with 1% SDS prior to sonication on a Bioruptor™ (Diagenode). The immunoprecipitation step involved incubation with different antibodies (Flag M2 beads (sigma, M8823), native NRPE1 antibody [61]) for at 4°C overnight. The antibody-chromatin complex was captured using protein Dynabeads A or G (Invitrogen) for 2 h, which were retained using a magnet. The beads were washed twice with high-salt buffer, low-salt buffer, LiCl buffer and TE buffer. Finally, the immunoprecipitated chromatin was reverse crosslinked and eluted from the beads with 10% (m/w) Chelex (Biorad) and digested with proteinase K at 50°C, followed by extraction of free DNA via ethanol precipitation. For the kinds of histone modification ChIP, non-crosslinked native ChIP [62] was performed. 14 days seedlings were harvested and ground to a fine powder in liquid nitrogen. Nuclei extraction was performed as described above. Resuspended nuclei in 500μl 1X MNase Buffer (50mM Tris-HCl pH 7.6, 5mM CaCl_2_, 0.1 mM PMSF, 1X protease inhibitor (Roche)) and incubated them with RNase A at 37°C for 30 min. Then total volume was made to 1ml with 1X MNase Buffer and added 4.5 μl Micrococcal Nuclease (NEB, M0247S) for incubation at 37°C for 30min. Stopped the reaction by adding EDTA (final 10 mM). Released nucleosomes by adding 0.1% SDS and rotated at 4°C for 3 hours. Then centrifuged at the maximum for 10min and diluted the supernatant with MNase dilution buffer (0.1% Triton X-100, 50 mM NaCl, 0.1mM PMSF and 1X protease inhibitor). In parallel, protein Dynabeads G (Invitrogen) beads should be incubated with antibodies (H3K9me2 Abcam, Ab1220), H3K4me3 (Millipore, 04-745), H3K36me3 (Abcam, Ab9050), H3 (ABclonal, A2348)) in ChIP Dilution Buffer (1.1% Triton X-100, 1.2 mM EDTA, 16.7 mM Tris-HCl pH 8.0, 167 mM NaCl, 1X protease inhibitor) at 4°C overnight. Beads-antibody complex was washed twice with 1ml MNase dilution buffer and added into diluted chromatins. The immunoprecipitation step was performed at 4°C for 5 hours. Then beads were washed twice with low salt wash buffer (50mM Tris-HCl pH 7.6, 10mM EDTA, 50mM NaCl, 0.1mM PMSF, 1X protease inhibitor), washed twice with middle salt wash buffer (50mM Tris-HCl pH 7.6, 10 mM EDTA, 100mM NaCl, 0.1mM PMSF, 1X protease inhibitor), washed once with high salt wash buffer (50 mM Tris-HCl pH 7.6, 10 mM EDTA, 150 mM NaCl, 0.1mM PMSF, 1X protease inhibitor), finally washed once with TE buffer (1 mM EDTA, 10 mM Tris-HCl pH8.0). Immune complexes were eluted by adding 200 μl elution buffer (0.1%SDS, 0.1M NaHCO_3_) at 65°C for 10 min, twice. Then treated with 2μl Proteinase K (10 mg/ml) for 1h at 45°C. Final DNA was recovered by Phenol/Chloroform extraction and ethanol precipitation.

For the Pol II Ser5p, total Pol II and Pol II Ser2p ChIP, method was modified from mNET-seq [63]. Briefly, crosslinked seedlings were ground to a fine powder in liquid nitrogen and nuclei was extracted with Honda Buffer as described above. Nuclei was performed with Micrococcal Nuclease treatment as described above for the fragmentation, then chromatin release was also performed as above. Added antibody-conjugated protein G beads (Active Motif 61085 for Pol II Ser5p, Active Motif for Pol II Ser2p and Ab817 for total Pol II) to the supernatant for 5-hours incubation at 4°C (antibody-beads conjugation should be done before: protein G beads were wash twice in NET-2 Buffer and incubated with antibody in 150 μl NET-2 Buffer rotating at 4°C overnight). After incubation, beads were washed 5 times in pre-cold NET-2 Buffer (50 mM Tris-HCl pH7.4, 150 mM NaCl, 0.05% NP-40). Elution and DNA recovery were performed as described above.

### pNET-seq and data analysis

pNET-seq was performed as described [41]. Briefly, 14-day old seedlings were harvested from 1/2MS media and were grounded in liquid nitrogen and resuspended in ice-cold lysis buffer (50mM HEPES pH 7.5, 150mM NaCl, 1mM EDTA pH 8.0, 1% Triton X-100, 10% glycerol, 5mM β-mercaptoethanol, 1mM PMSF, 2μg/μl Aprotinin and 2μg/μl Pepstatin A) for 30 mins on ice. After lysis mixture was filtered and centrifuged to discard the supernatant, Nuclear pellet was washed once with ice-cold HBB (25mM Tris-HCl pH7.6, 0.44M sucrose, 10mM MgCl_2_, 0.10% Triton-100, 10mM β-mercaptoethanol) and HBC (20mM Tris-HCl pH7.6, 0.352M sucrose, 8mM MgCl_2_, 0.08% Triton-100, 8mM β-mercaptoethanol and 20% glycerol) buffers, respectively. Washed nuclear pellet was resuspended in MNase reaction buffer (20mM Tris-HCl pH 8.0, 5mM NaCl, and 2.5mM CaCl_2_, 20U MNase (TAKARA)). The reaction mixture was incubated at 37°C for 5 minutes and the reaction was later stopped with final con. 0.02M EDTA. Then sonication was performed 1s ON/OFF for 10 cycles to release chromatin and centrifuged to obtain the supernatant. The collected supernatant was diluted 5-8 times with ice-cold NET-2 buffer (50mM Tris-HCl pH7.4, 150mM NaCl, and 0.05% NP-40). The diluted supernatant was incubated with Pol II Ser5p antibody (MBL Life science, MABI0603) coated Dynabeads (Invitrogen, 11201D) for 2 hours at 4°C and the beads containing complex was washed 8 times with ice-cold NET-2 buffer. Finally, beads were resuspended with PNK reaction mixture containing 75 μl PNK buffer, 10 μl PNK enzymes (NEB) and 15 μl of 10mM ATP and incubated at 37°C for 10 mins on a Thermomixture (1400 rpm). After that supernatant was removed and beads were washed once with NET-2 buffer. Then beads were followed RNA extraction with Trizol reagent. The extracted RNA was performed gel size selection with 8% PAGE gel and the gel corresponding to 35-100bp size of RNA was collected. After size-selected RNA was recovery from the gel, NEXTflex TM small RNA-Seq Kit V3 was used to construct the small RNA libraries. Finally, the libraries were recovered from the 6% PAGE gel corresponding to 140-250 bp size and undergone high throughput deep sequencing. For the pNET-seq data analysis, only R2 of paired-end reads were trimmed for Illumina adaptors and then the reads <10bp were filtered by using the Cutadapt (v1.9.1). Filtered reads were aligned to the Arabidopsis reference genome (TAIR10) using Bowtie 2 version 2.3.0 with default settings. BAM files were converted to bigWig using RPGC normalization method in deepTools v2.26.0. MACS2 was used to identify differently regulated Ser5 peaks in *nrpe1* vs Col-0. bigWig files were used for visualization in IGV and calculating rawcounts in indicated regions with deepTools. deepTools were also used for correlation plot and Spearman’s correlation coefficients calculation between two replicates using 100bp bins. pNET-seq data have been deposited in the Gene Expression Omnibus database (accession no. GSE125096). Other published data are obtained for analysis in this study: H3K4me2 and H3K4me3 ChIP-seq (GSE49090), H3K9me2 ChIP-seq (GSE118063), whole genome bisulfite sequencing (GSE80303), NRPE1 RIP-seq (GSE70290), anti NRPE1 ChIP-seq (GSE61192).

## Acknowledgements

This work was supported by grants from the National Key R&D Program (2016YFA0500800 to The Sun Lab), the National Natural Science Foundation of China (No.31571258 to Liangyu Liu, and No.31571322 and 91740105 to The Sun Lab), the Youth Innovative Research Team Project of Capital Normal University (to Liangyu Liu). The Sun Lab was supported by Tsinghua University Initiative Scientific Research Program, Tsinghua-Peking Joint Center for Life Sciences, and the 1000 Young Talent Program of China. W.X. was supported by postdoctoral fellowship from Tsinghua-Peking Joint Center for Life Sciences. We thank all The Sun Lab members for the useful discussions and suggestions, Dr. Yijun Qi for the *nrpd1a-3, rdr2-1, dcl3-1, ktf1, drd1* and *rdr6* mutants, Dr. Xiaofeng Fang for the help of siRNA blotting and Pol V ChIP analysis, and Dr. Zhicheng Dong for the help of pNET-seq.

## Author contributions

Q Sun and J Zhou designed the project, Q Sun, J Zhou and L Liu carried out experiments and analyzed the data; J Zhou and Q Li generated the plant materials; W Xu desined the primers for bisulfite sequencing on FLC, and K Li, W Xu and ZW Wang assisted the data analysis. All authors analyzed the data together, and J Zhou, L Liu and Q Sun wrote the paper with the input from all other authors.

---

**Figure S1.**
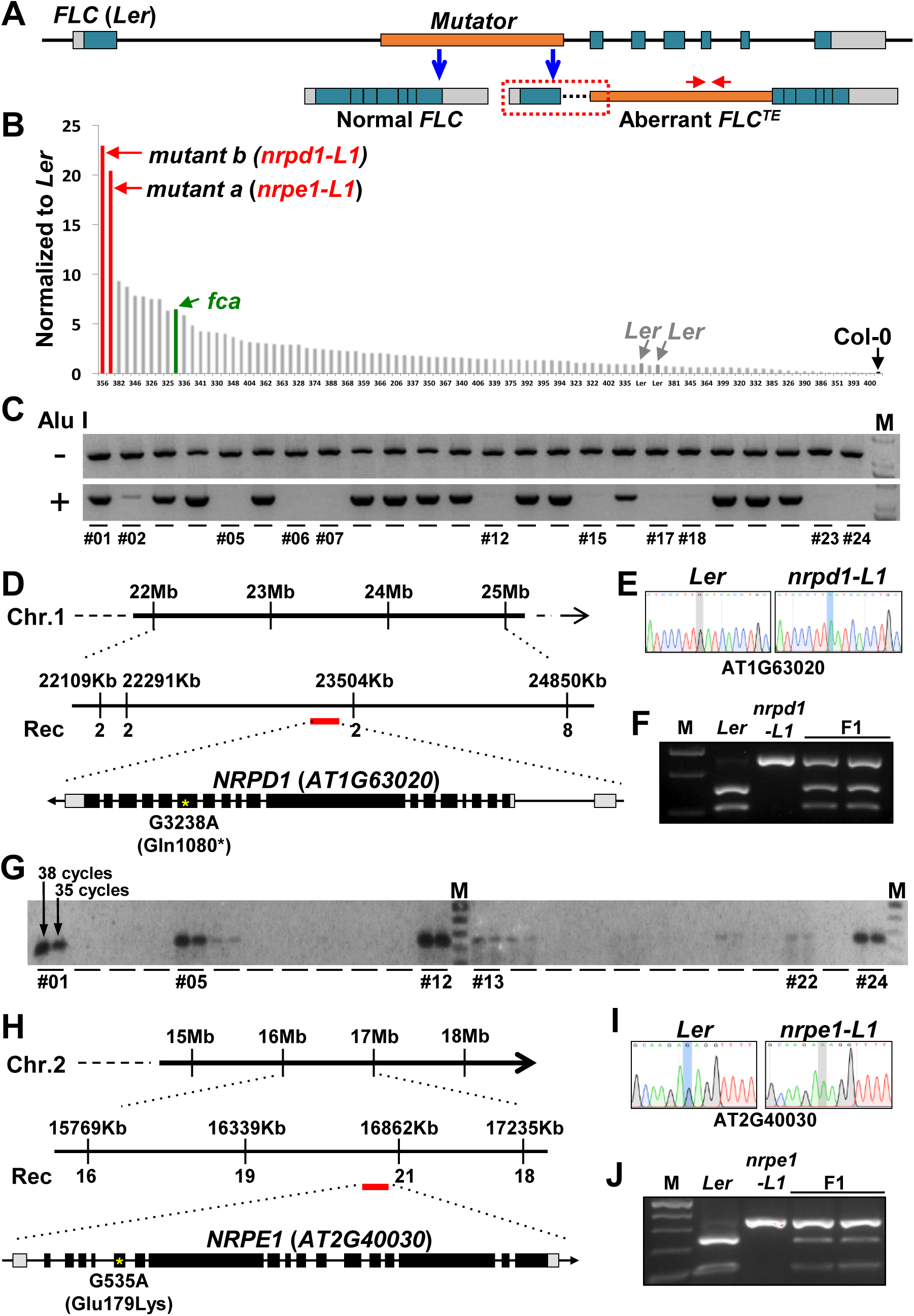
Identifying *NRPD1* and *NRPE1* as the repressors of aberrant transcription of *FLC*^*TE*^. (**A**) Schematic map of the *Ler FLC* locus, including a Mutator-like TE (orange box). Normal *FLC* and TE-mediated aberrant *FLC*^*TE*^ are listed below the whole locus. Red dotted box indicates unidentified 5’ sequence of aberrant *FLC*^*TE*^ according to published data [25]. Gray and blue boxes indicate UTR region and exons, respectively. (**B**) Relative expression levels of Mutator-like TE from EMS-generated mutants are normalized to *Ler* control as revealed by RT-qPCR. Primers used for qPCR are shown in Figure 1A (red arrows) and supplemental table 1. (**C**) Chop-PCR based selection of F2 mapping population. After F2 plants (#2-23) genomic DNA digestion with a methylation-sensitive restriction enzyme Alu I, 31 cycles PCR products were analyzed. The selected recombinants (#02, #05, #06, #07, #12, #15, #17, #18 and #23), which were then used for Map-based cloning, have the same PCR signal as the original mutant (#24) but significantly reduced signal compared to *Ler* (#01). (**D and H**) Mapping and isolation of the *NRPD1* and *NRPE1* gene locus. The point mutation is highlighted by a yellow asterisk and results in a premature stop codon in *nrpd1-L1* mutant and an amino acid transition from glutamic acid to lysine in *nrpe1-L1* mutant. (**E and I**) Sequencing chromatograms show both G-to-A transitions in the *AT1G63020* (*NRPD1*) and the *AT2G40030* (*NRPE1*) genes sequence. (**F and J**) Genotyping of plants was based on the polymorphism of Bcl I-digesting products (for mutant *nrpd1-L1*) and MnI I-digesting products (for mutant *nrpe1-L1*). (**G**) RT-PCR based methods to select the recombinants in a segregated F2 population (crossed with Col-0) for fine mapping of *nrpe1-L1*. Results of a typical analysis of F2 plants (#1–24) by RT-PCR for 35 and 38 cycles. Map-based cloning was then applied to a selected recombinant population (#01, #05, #12, #13 and #22); the original mutant #24, was used as a positive control.

**Figure S2.**
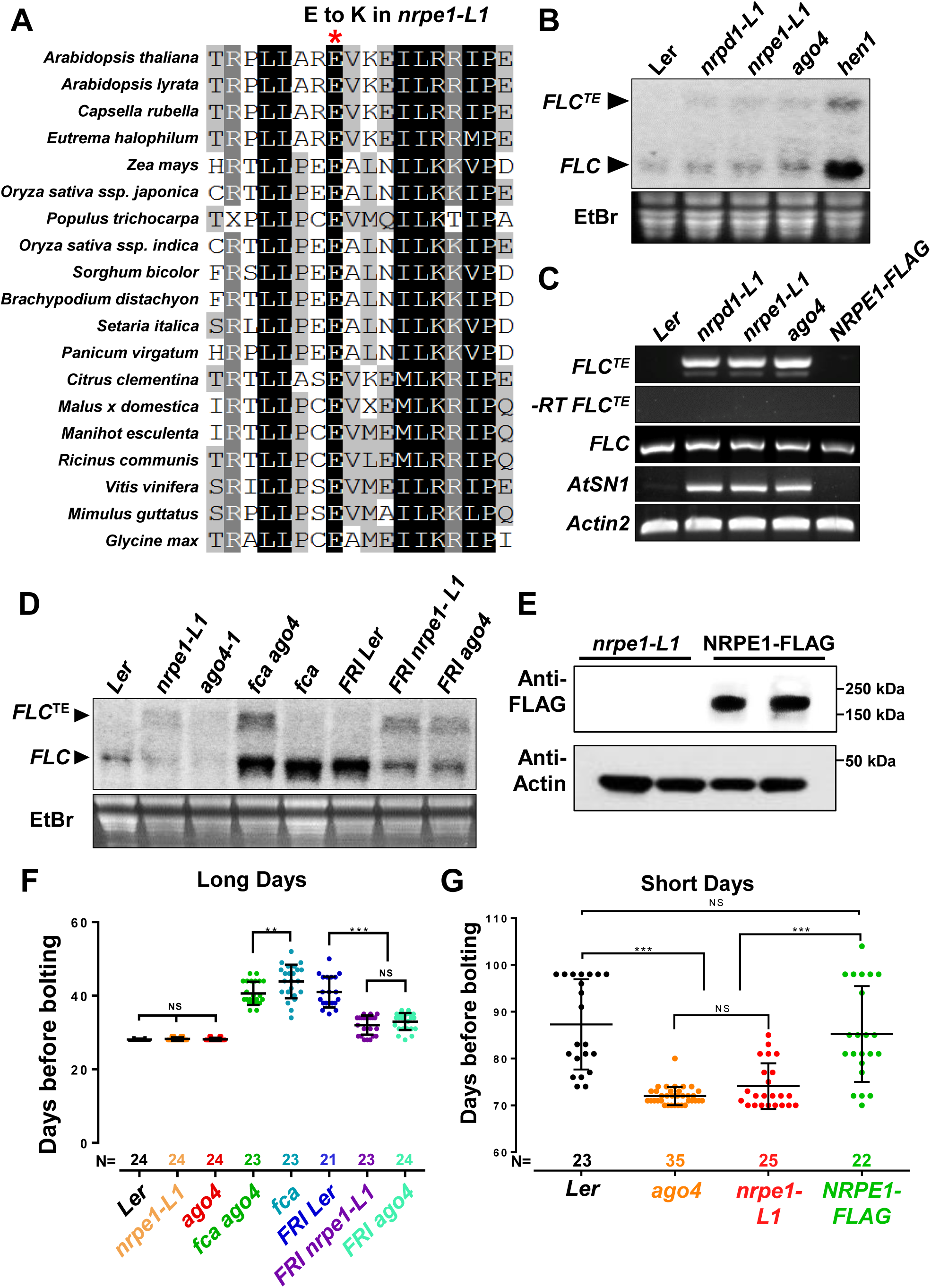
*nrpe1* mutant modulates flowering. (**A**) The mutated residue glutamic acid (E, marked by a red asterisk) is located in a conserved NRPE1 region in plants. (**B**) Northern blot analysis shows both *nrpd1-L1* and *nrpe1-L1* mutants have the same aberrant *FLC*^*TE*^ transcript as *ago4* and *hen1* mutants. Total RNA was stained with EtBr as a loading control. (**C**) ssRT-PCR results indicate aberrant *FLC*^*TE*^ in the mutants and complementation line *pNRPE1::NRPE1:FLAG/nrpe1-L1.* And RNA transcript from Pol V-repressed target *AtSN1* is de-repressed in *nrpd1-L1, nrpe1-L1* and *ago4. Actin2* amplicons were used as internal controls. Samples without reverse transcription amplified by *FLC*^*TE*^ primers was used as negative controls. (**D**) Aberrant *FLC* transcription is independent of FRI and FCA. *FLC*^*TE*^ was detected by northern blot analysis. Total RNA was stained with EtBr as a loading control. (**E**) FLAG-tagged NRPE1 is enriched in the complementation line, as revealed by immunoblotting. Actin protein was used as a loading control. (**F**) Flowering time (number of days from germination to bolting) were measured in greenhouse under long days (LDs, 16-hours light / 8-hours dark). Data are represented as mean±s.d., and more than 20 individual plants for each genetic material were used for analysis. Asterisks denote statistical significance of mean values from wild type (* p<0.05, **p<0.01, and ***p<0.001). (**G**) Flowering time was measured based on number of days from germination to bolting. Plants were grown in greenhouse under short-day conditions (8-hour light / 16-hour dark). Data are represented as mean±s.d.. The calculation was based on ⩾20 independent plants, ***p < 0.001.

**Figure S3.**
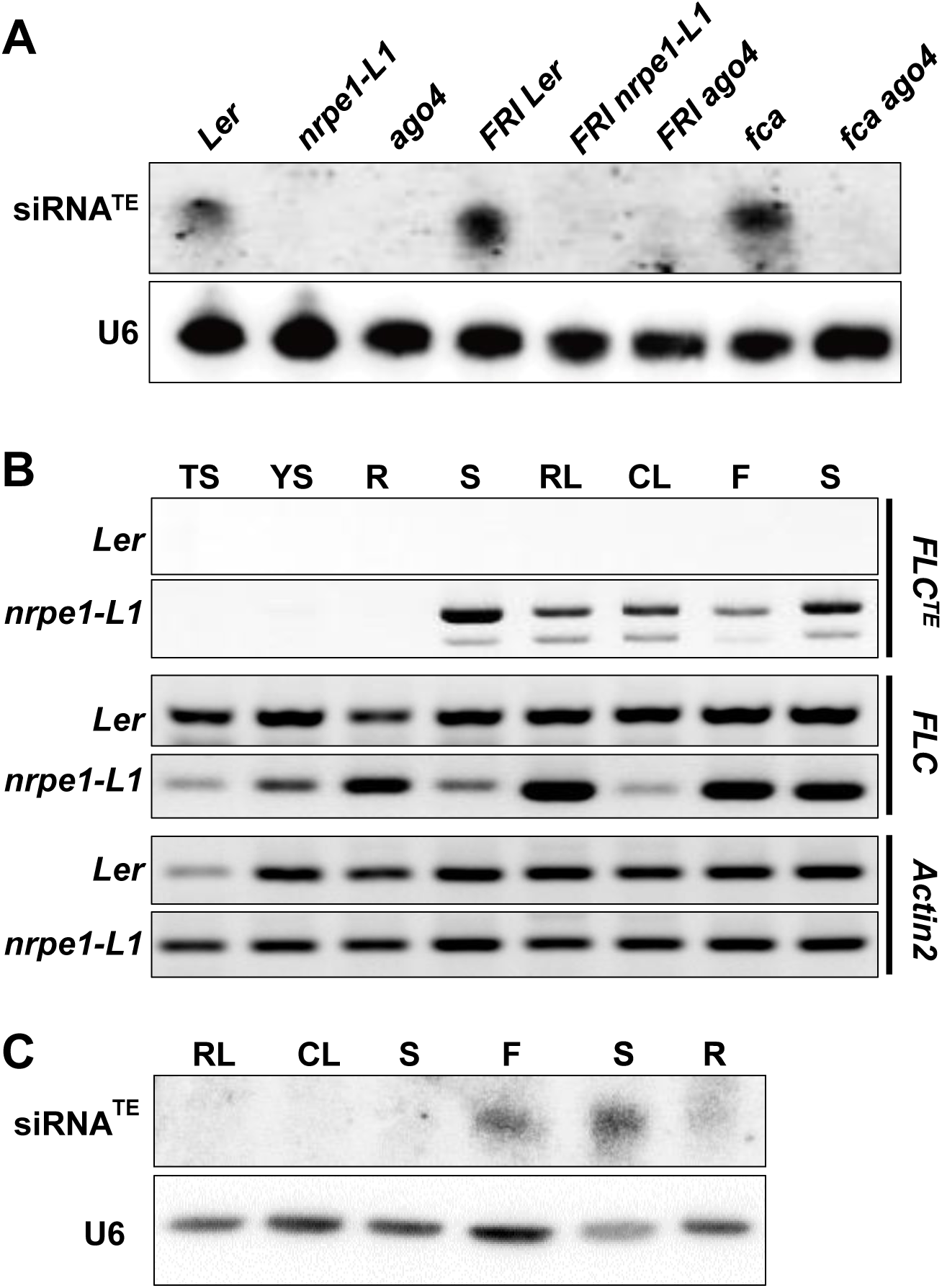
RdDM mutants reduce siRNA^TE^ accumulation but repression of *FLC* cryptic initiation is independent of local siRNA biogenesis. (**A**) Northern blot analysis shows the levels of siRNA^TE^ in wild type *Ler* and mutants. *U6* RNA was used as a loading control. (**B**) Profiling of *FLC*^*TE*^, *FLC* and *Actin2* in *Ler* and *nrpe1-L1* by RT-PCR and ssRT-PCR. (**C**) Profiling of siRNA^TE^ and *U6* RNA in *Ler* by northern blot analysis. TS: total siliques; YS: yong siliques; R: roots; S: stems; RL: rosette leaves; CL: cauline leaves; F: flowers; S: seedlings.

**Figure S4.**
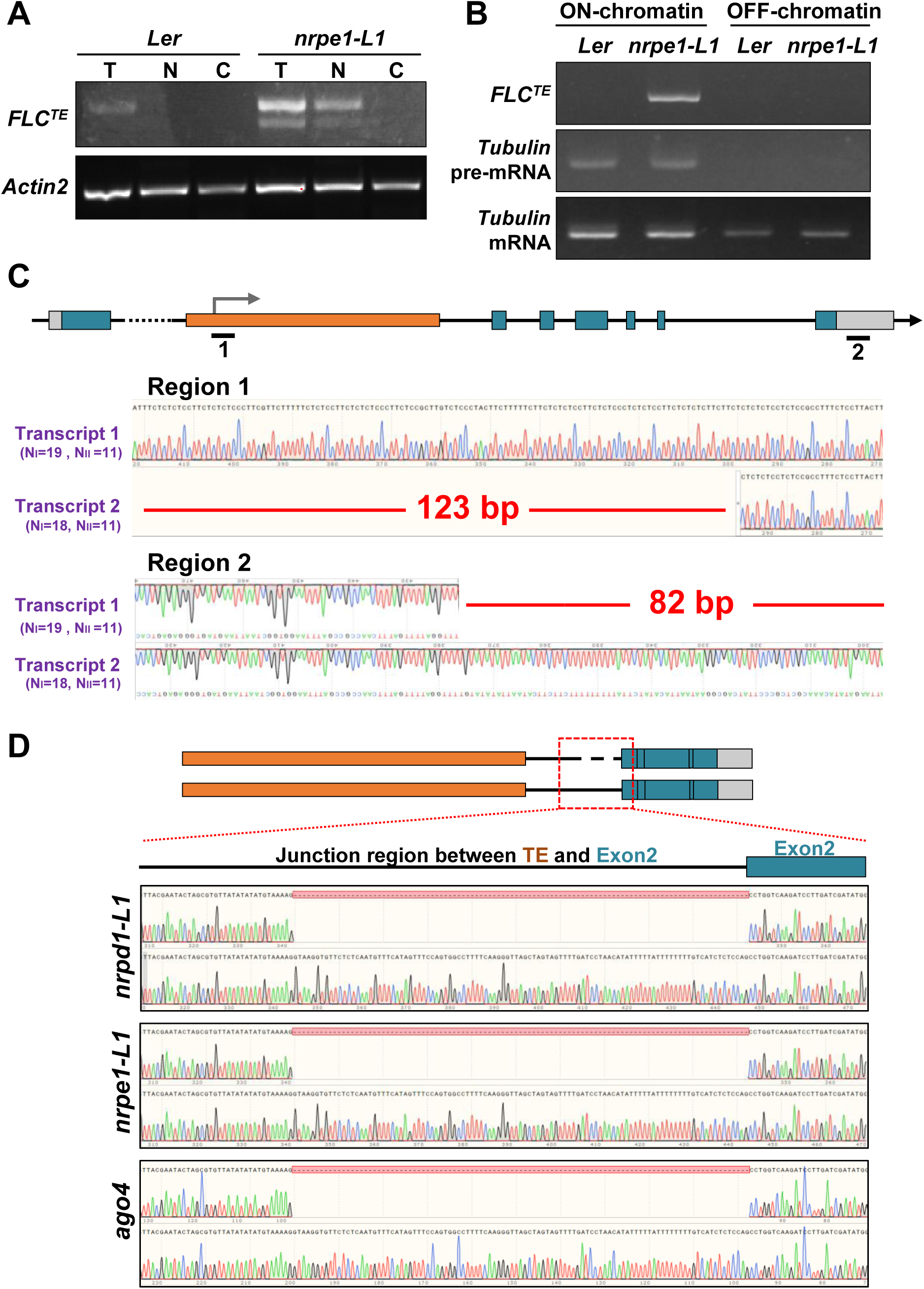
Transcription initiation sites, termination sites and splicing sites of *FLC*^*TE*^. (**A**) *FLC*^*TE*^ transcripts measured by ssRT-PCR are enriched in *nrpe1-L1* compared to *Ler*, especially in isolated nuclei (N) and total cells (T), but not in cytoplasm (C). (**B**) *FLC*^*TE*^ transcripts are associated with chromatin and are hardly detected in the off-chromatin fraction. *Tubulin* pre-mRNA was used as control. (**C**) Sequencing chromatograms (clones from **Figure 2B** and **2C**) show *FLC*^*TE*^ has two transcription initiation sites (region 1) and two termination sites (region 2). (**D**) Sequencing results of **Figure S2C** *FLC*^*TE*^ bands from *nrpe1-L1, nrpd1-L1* and *ago4* show two splicing isoforms at the intron region upstream of exon2.

**Figure S5.**
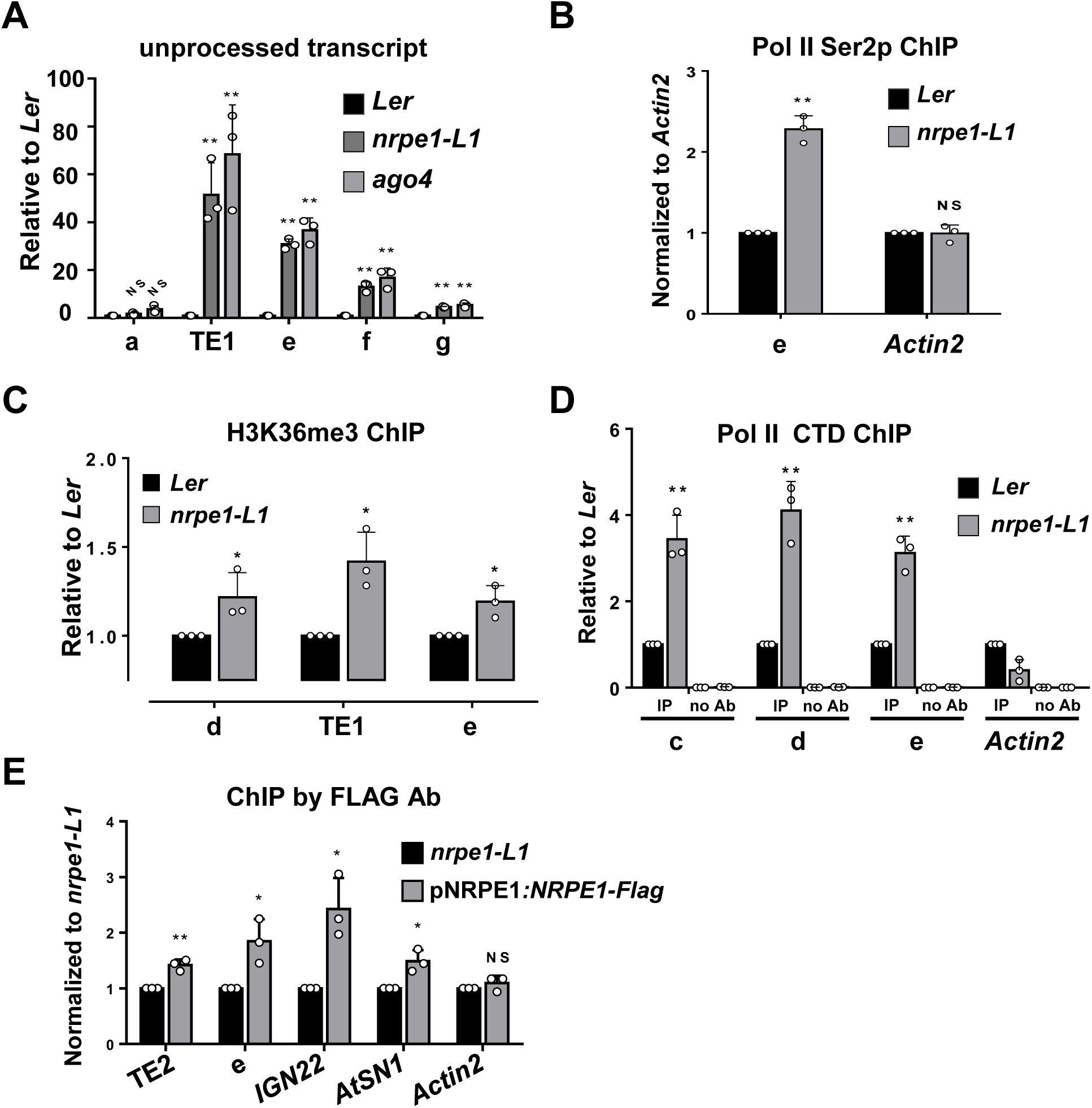
NRPE1 inhibits the transcription of *FLC*. (**A**) RT-qPCR shows the increased levels of unprocessed *FLC* RNA transcripts in *nrpe1-L1* and *ago4*. In **C**-**G**, asterisks denote statistical significance of mean values from wild type (*p<0.05 and **p<0.01). (**B and D**) Pol II ChIP-qPCR analysis shows binding of total Pol II (Pol II CTD) and Elongated Pol II (Pol II Ser2) on *FLC* gene between *Ler* and *nrpe1-L1*, respectively. Error bars represent s.d. (n=3). (**C**) H3K36me3 ChIP-qPCR data indicates increased active histone marks on *nrpe1-L1 FLC* gene. (**E**) FLAG antibody against NRPE1:FLAG driven by its native promoter in the *nrpe1-L1* background, with *nrpe1-L1* used as the control line. Values were normalized to input and control line signals. *IGN22* and At*SN1* were used as positive controls and *Actin2* was used as a negative control. Error bars, s.d., were calculated based on values from 3 representative experiments.

**Figure S6.**
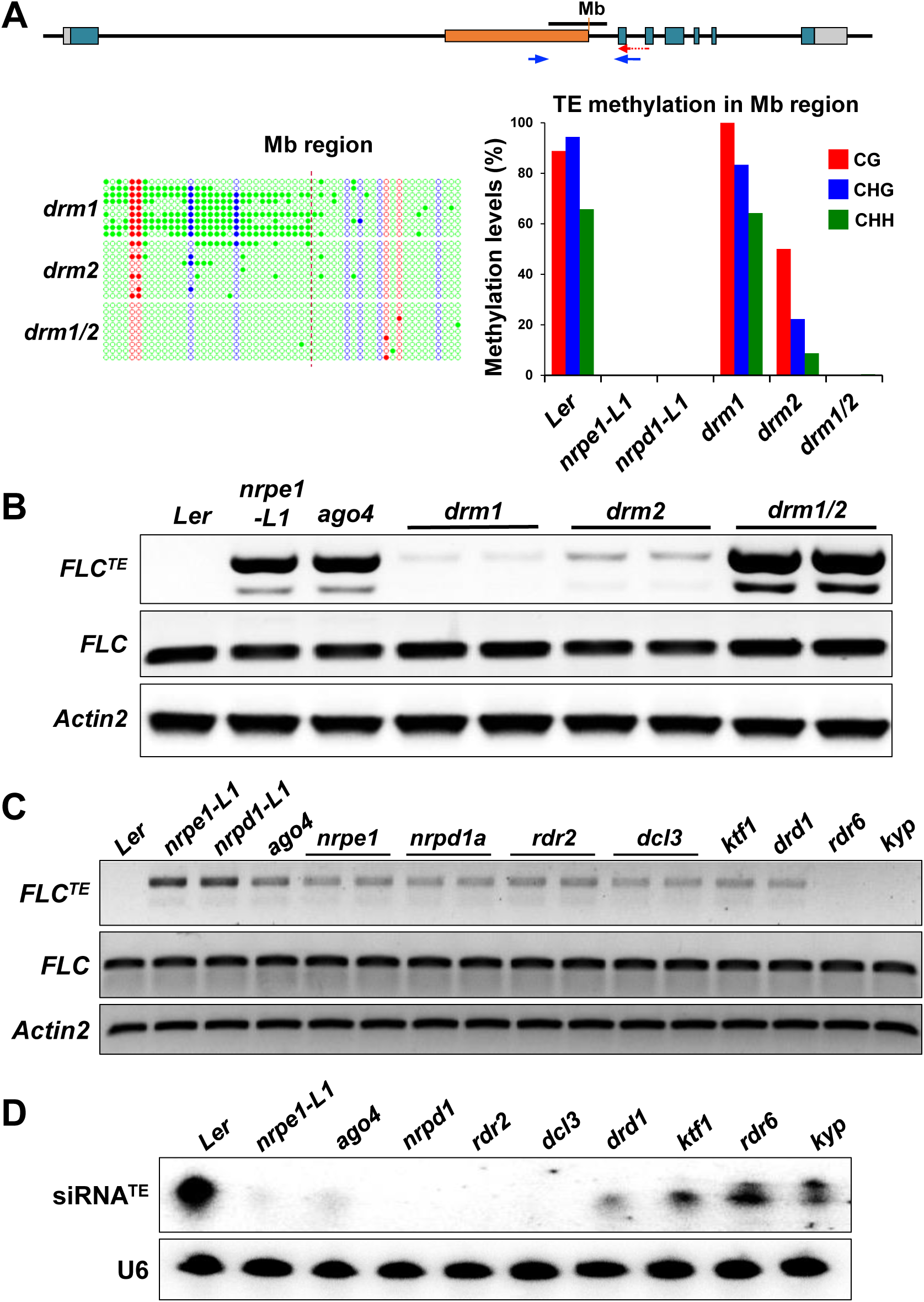
Repression of *FLC* cryptic initiation relies on RdDM-mediated DNA methylation but not local siRNA biogenesis. (**A**) DNA methylation of *FLC* TE is regulated by DRM1 and DRM2. DNA methylation on the sense strand of *FLC* (region Mb) was analyzed by bisulfite sequencing. Red dashed line marks the right border of the TE region. CG, CGH and CHH methylation levels (%) in region Mb are shown on the right. (**B**) CHH DNA methyltransferases DRM1 and DRM2 are both required to repress *FLC* cryptic transcription initiation. (**C**) Major components of the RdDM pathway suppress *FLC* cryptic initiation. *FLC*^*TE*^ and *FLC* levels were measured by ssRT-PCR in mutants including *nrpe1-L1, nrpd1-L1, ago4, nrpe1, nrpd1, rdr2, dcl3, ktf1, drd1, rdr6* and *kyp*. Here, except the *Ler* ecotypes of *nrpe1-L1, nrpd1-L1* and *ago4* mutants, the others were obtained by crossing the Col-0 ecotype mutant with *Ler*, and the independent crossing lines carrying *FLC-TE* locus were isolated. *Actin2* was used as an internal control. Red and blue arrowheads shown in Figure S5A indicate primers for ssRT and PCR respectively. (**D**) siRNA^TE^ levels were barely detected in *nrpe1-L1, ago4, nrpd1a, rdr2* and *dcl3*, dramatically reduced in *drd1*, and not altered in *ktf1, rdr6* and *kyp. U6* RNA was used as a loading control for northern blot analysis.

**Figure S7.**
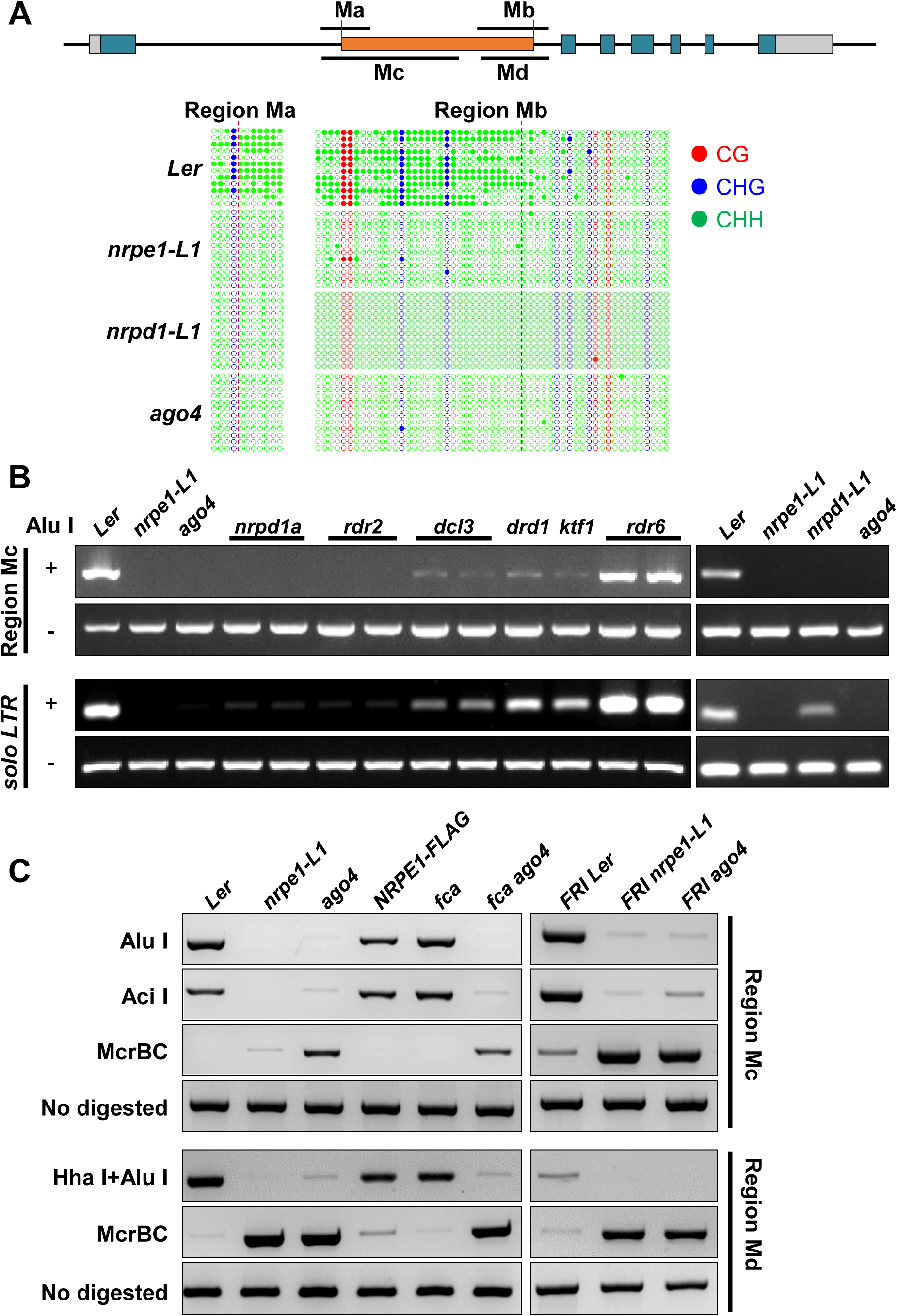
RdDM pathway mutants reduce DNA methylation on *FLC*. (**A**) Ma and Mb as sense regions are used for DNA methylation analysis respectively, and dashed black lines indicate the Mutator-like transposon borders. CG, CHG and CHH DNA methylation levels are dramatically reduced in *nrpe1, nrpd1* and *ago4*, as measured by bisulfite sequencing of regions Ma and Mb. (**B**) Major RdDM mutants including *nrpe1-L1, nrpd1-L1, ago4, nrpd1, rdr2, dcl3, ktf1* and *drd1* are depleted of CHH DNA methylation on region Mc, as revealed by Chop-PCR. The *solo LTR* was used as a control. (**C**) DNA methylation levels on region Mc and Md assayed by Chop-PCR are independent of FRI and FCA but are rescued by introducing *pNRPE1::NRPE1:FLAG*. Undigested DNA and McrBC (recognizes methylated DNA)-digested DNA were used as controls.

**Figure S8.**
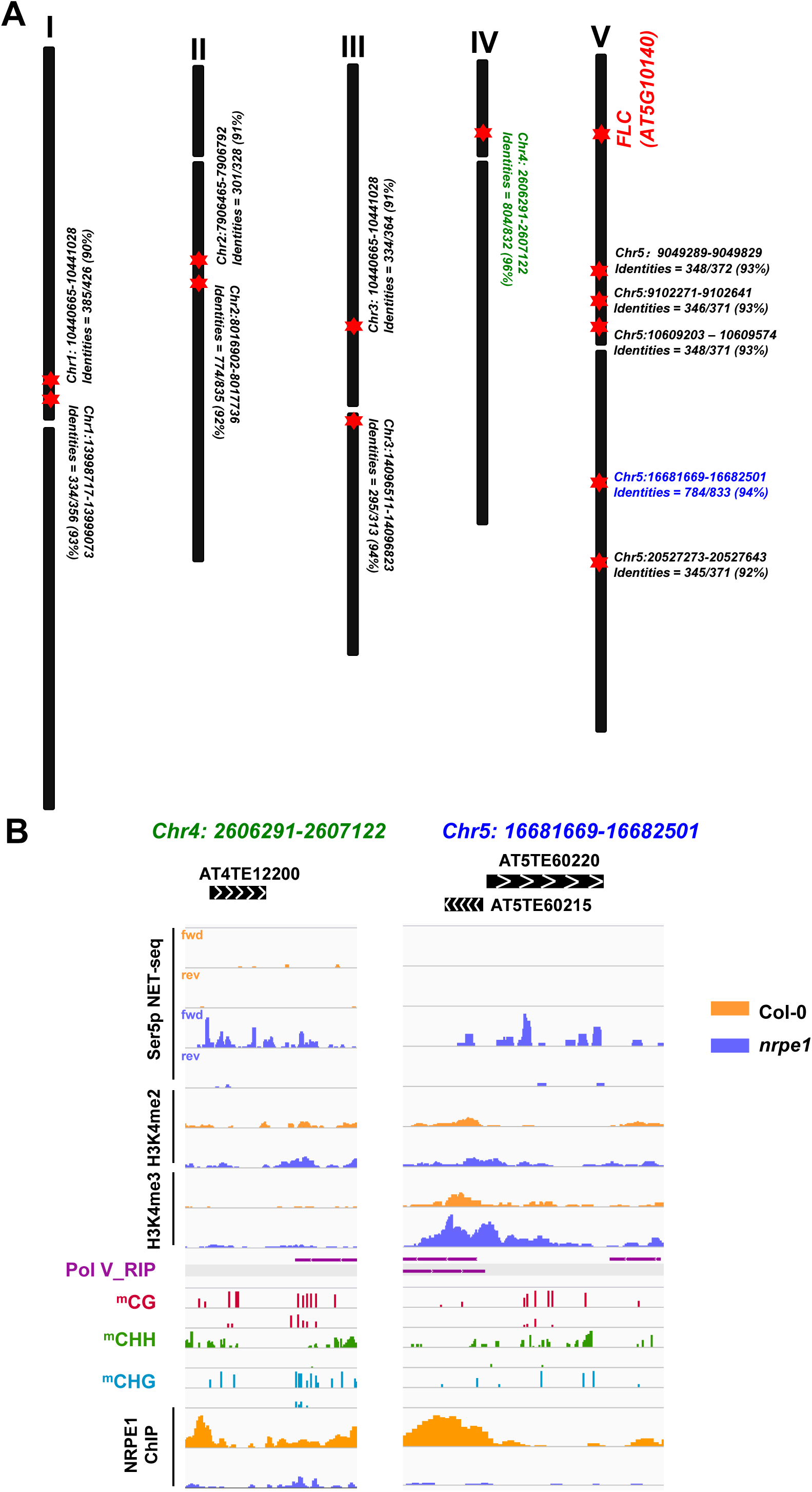
NRPE1 is associated with *FLC-TE* homologous loci. (**A**) Distribution of *FLC-TE* homologous loci in Col-0 genome. (**B**) Snapshot of two *FLC-TE* homologous loci (marked in green and blue in Figure S8A) with indicated different modifications from IGV.

**Figure S9.**
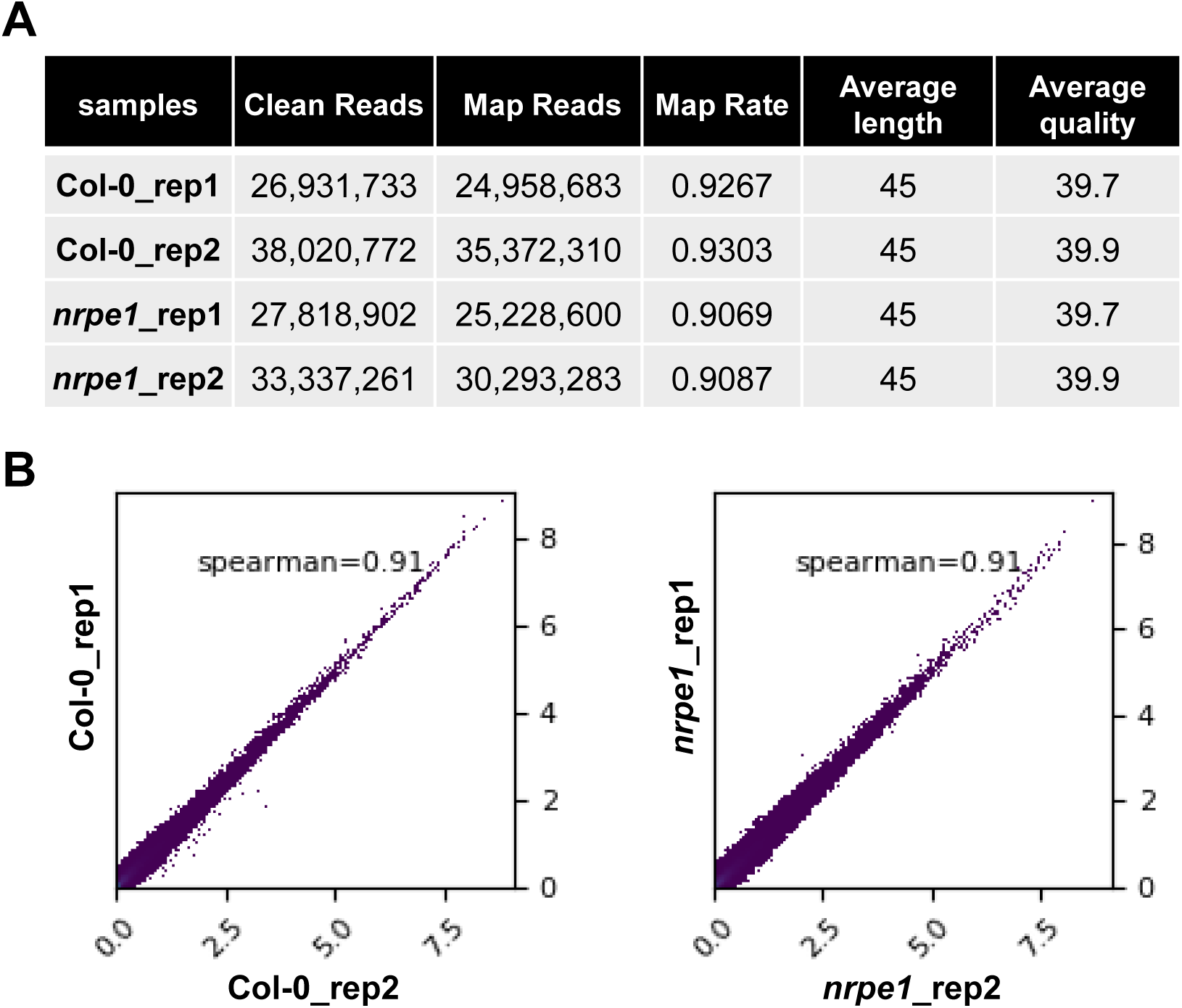
Assessment of the the Pol II Ser5p NET-seq data. (**A**) Basic information of NET-seq data of two replicates of indicated samples. (**B**) Scatterplots of two replicates of indicated samples. Values are calculated per bin with 100bp. Spearman’s correlation coefficients were indicated.

**Figure S10.**
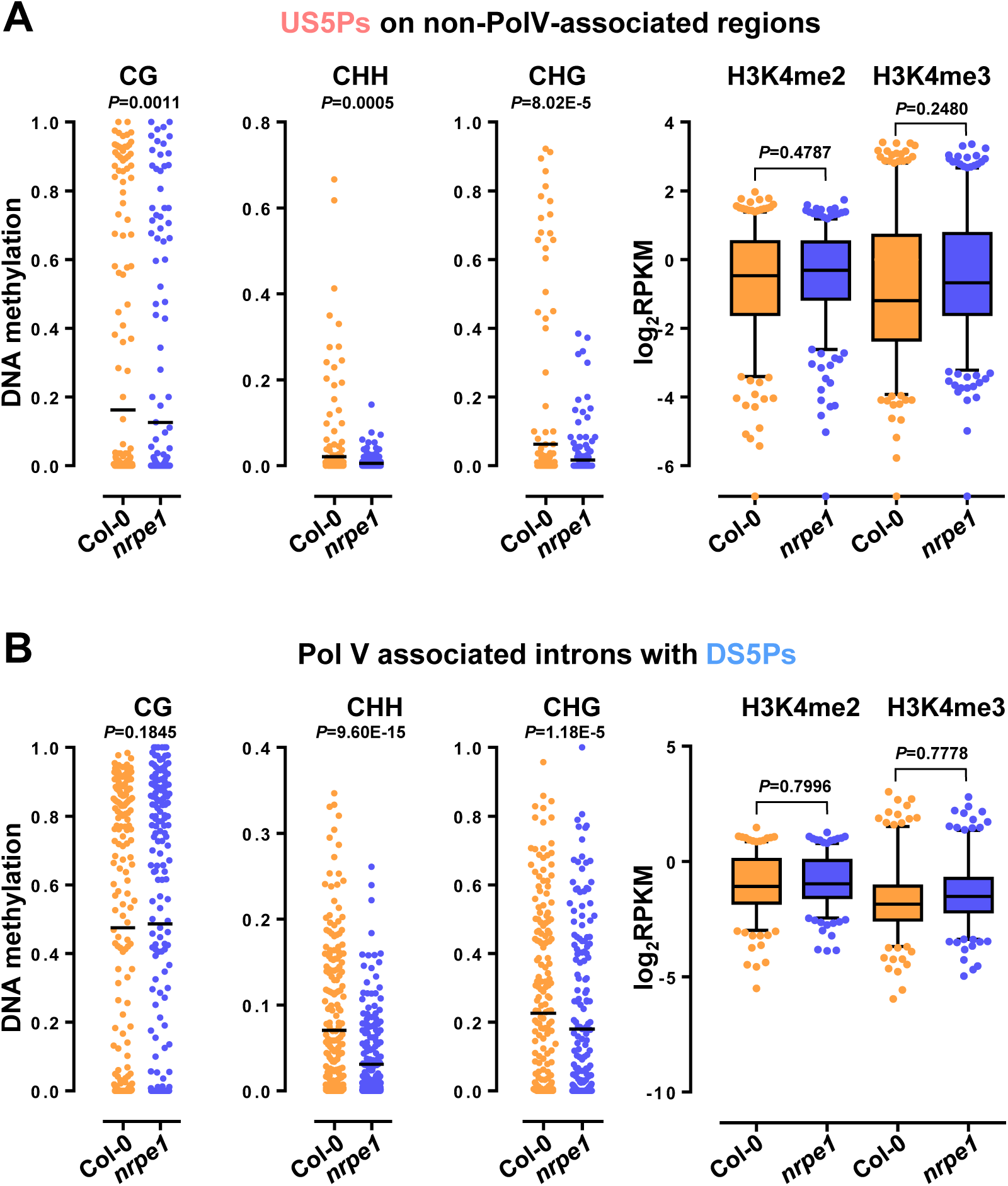
The levels of DNA methylation and H3K4me2/3 modifications in differentially modulated Ser5p regions. (**A**) DNA methylation levels and H3K4me2/3 levels of US5Ps on non-Pol V associated regions. (**B**) DNA methylation levels and H3K4me2/3 levels of downregulated Pol V associated-introns according to Ser5p pNET-seq data.

**Figure S11.**
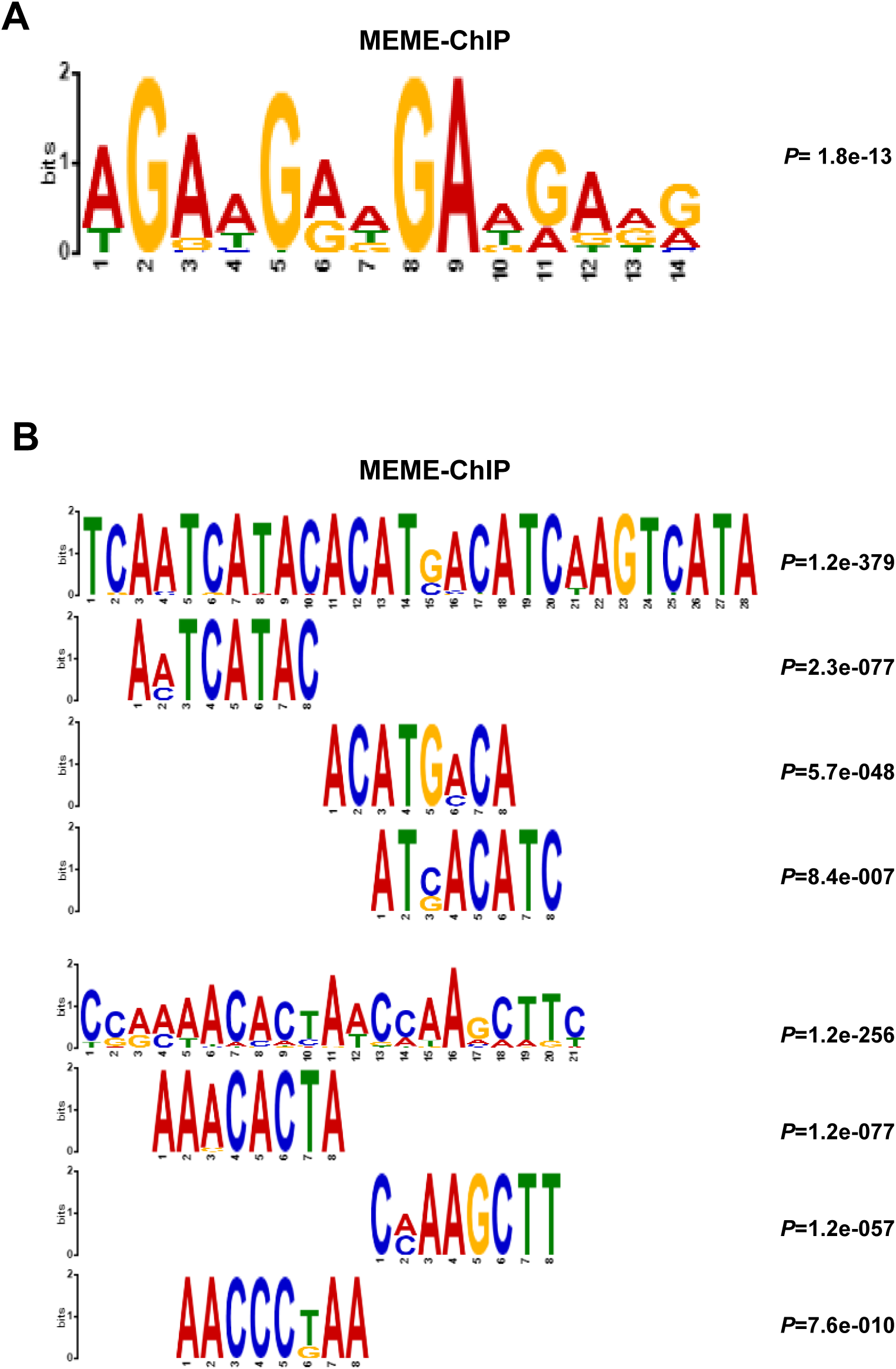
Motifs analysis. (**A**) Motif analysis of Pol V associated introns with US5Ps in *nrpe1* versus Col-0 using online software MEME-ChIP. (**B**) Motif analysis of Pol V associated regions (Pol V RIP-seq data) using online software MEME-ChIP.

**Table S1.**
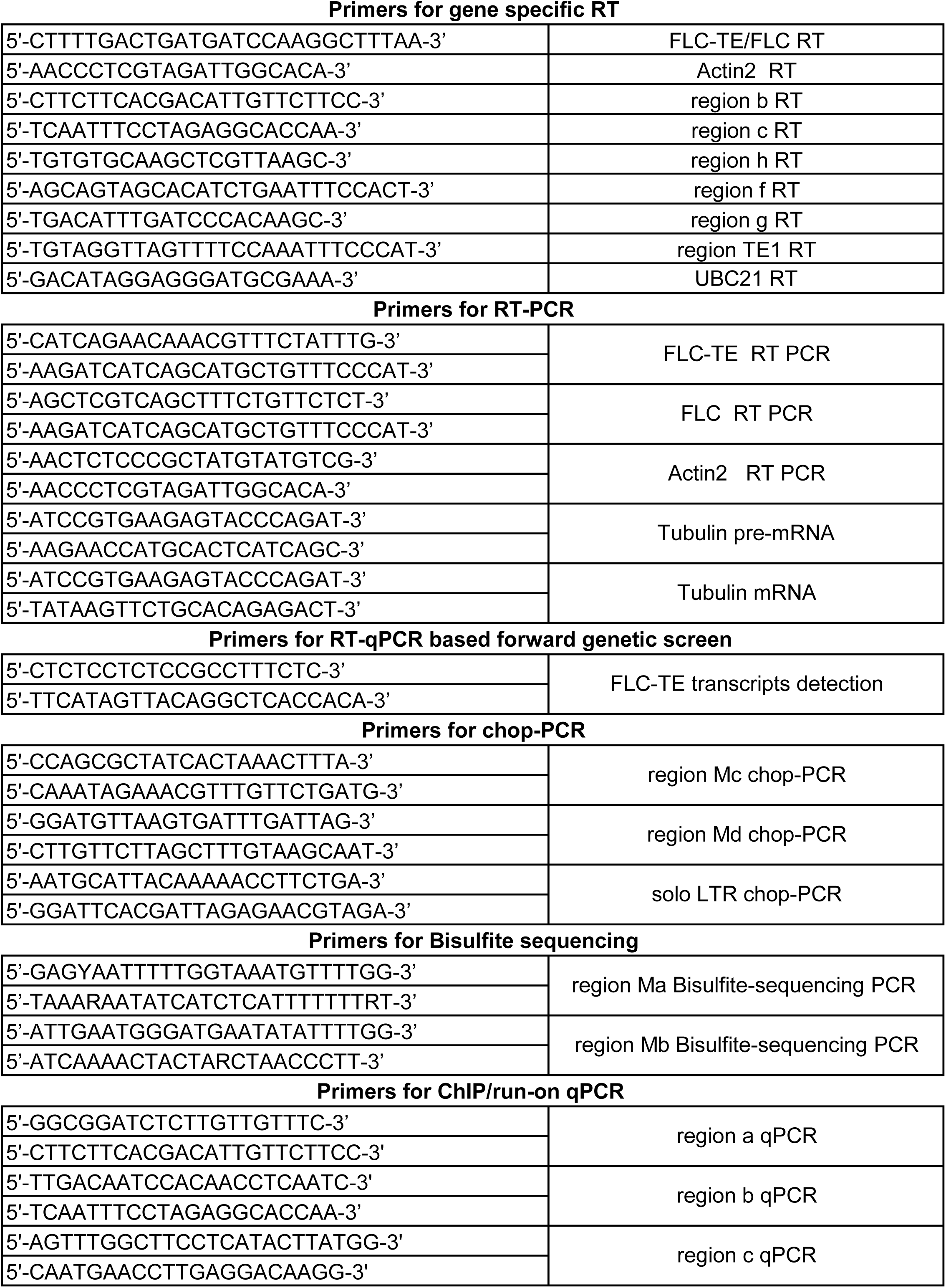
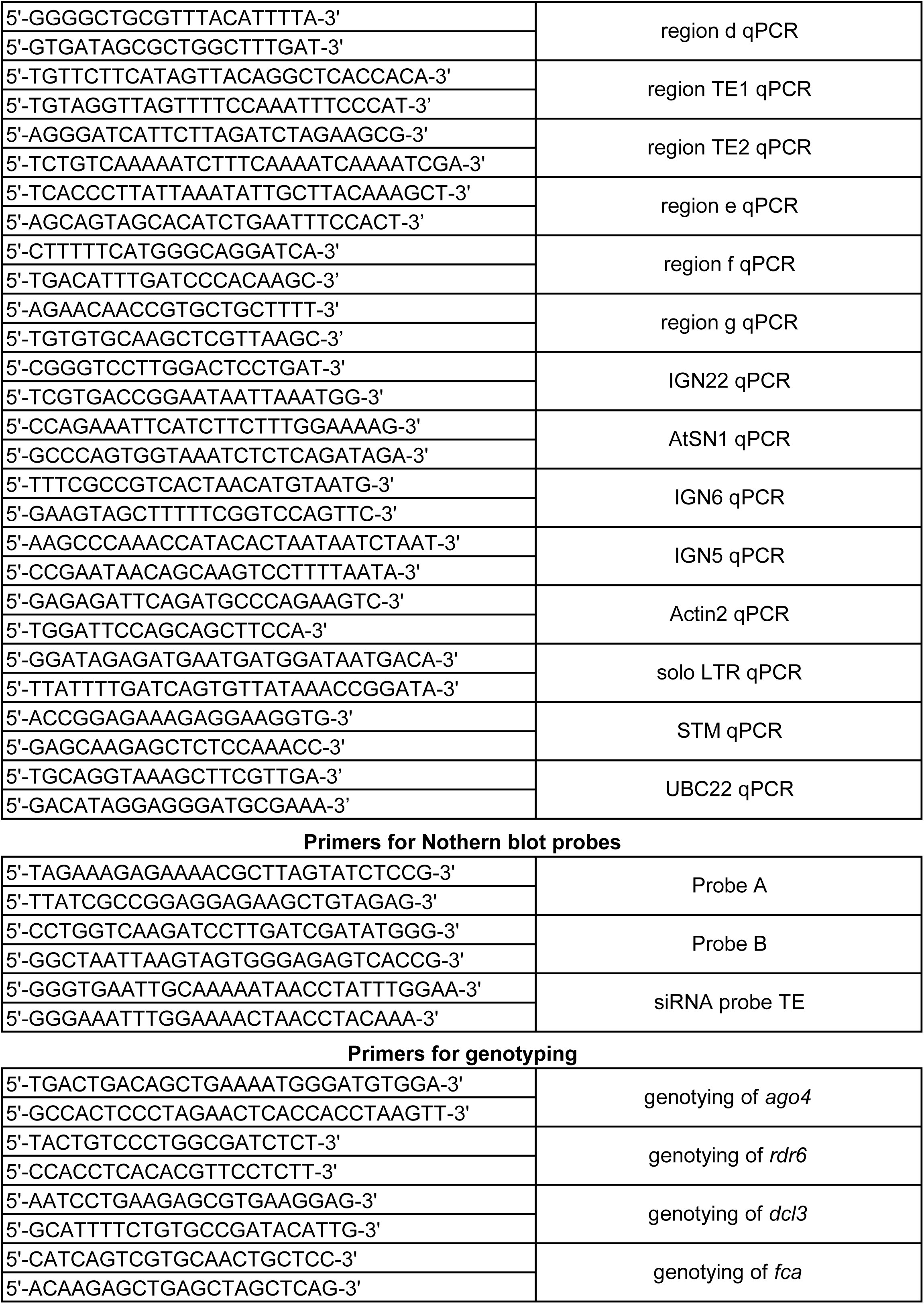
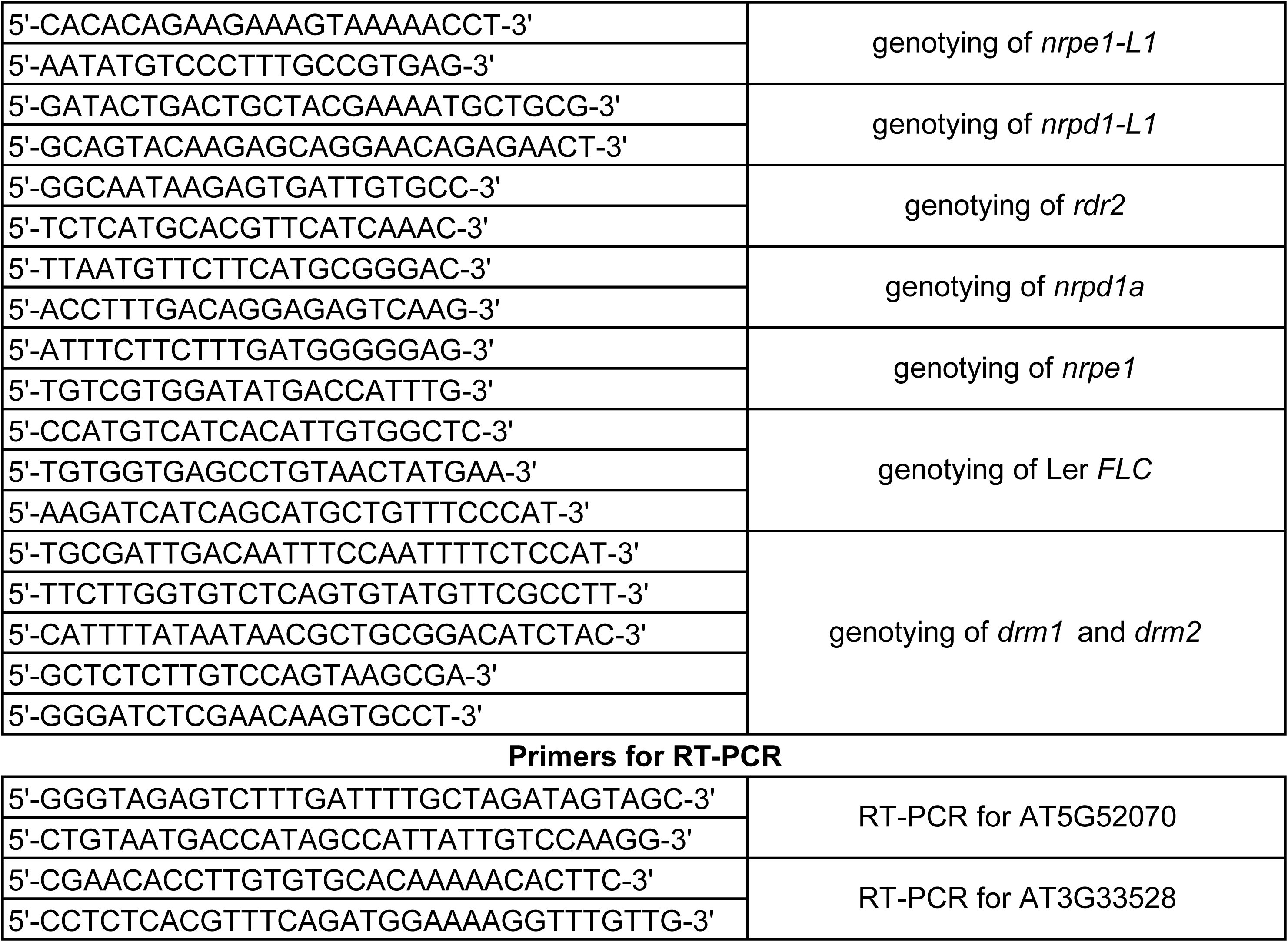
List of primers used in this study.

**Table S2.** Pol V associated introns with US5Ps.

| introns |  |  |  |  | intronic TEs |  |  |  |  |  |
| --- | --- | --- | --- | --- | --- | --- | --- | --- | --- | --- |
| chr. | intron_start | intron_end | introns | strand | chr. | TE_start | TE_end | intronic TEs | TE_subfamily | strand |
| Chr1 | 702738 | 702827 | AT1G03030 intron7 | + |  |  |  |  |  |  |
| Chr1 | 703099 | 703194 | AT1G03030 intron9 | + |  |  |  |  |  |  |
| Chr1 | 703382 | 703481 | AT1G03030 intron11 | + |  |  |  |  |  |  |
| Chr1 | 1989527 | 1989667 | AT1G06500 intron1 | + |  |  |  |  |  |  |
| Chr1 | 2051397 | 2051483 | AT1G06690 intron7 | - |  |  |  |  |  |  |
| Chr1 | 2051525 | 2051601 | AT1G06690 intron8 | - |  |  |  |  |  |  |
| Chr1 | 2051694 | 2051798 | AT1G06690 intron9 | - |  |  |  |  |  |  |
| Chr1 | 2398690 | 2399098 | AT1G07740 intron1 | - |  |  |  |  |  |  |
| Chr1 | 4967265 | 4967343 | AT1G14518 intron1 | - |  |  |  |  |  |  |
| Chr1 | 6709784 | 6709888 | AT1G19394 intron1 | + |  |  |  |  |  |  |
| Chr1 | 7376508 | 7376588 | AT1G21070 intron1 | - |  |  |  |  |  |  |
| Chr1 | 8644091 | 8644161 | AT1G24370 intron1 | - |  |  |  |  |  |  |
| Chr1 | 8644091 | 8644161 | AT1G24370 intron1 | - |  |  |  |  |  |  |
| Chr1 | 8771871 | 8771957 | AT1G24807 intron1 | - |  |  |  |  |  |  |
| Chr1 | 8772829 | 8773427 | AT1G24807 intron6 | - |  |  |  |  |  |  |
| Chr1 | 8776340 | 8781949 | AT1G24880 intron3 | + | Chr1 | 8778395 | 8778654 | AT1TE28320 | TAT1 ATH | + |
| Chr1 | 8782063 | 8782163 | AT1G24880 intron4 | + |  |  |  |  |  |  |
| Chr1 | 8798868 | 8804477 | AT1G25054 intron3 | + | Chr1 | 8800897 | 8801182 | AT1TE28395 | TAT1 ATH | + |
| Chr1 | 8812202 | 8813944 | AT1G25097 intron2 | - |  |  |  |  |  |  |
| Chr1 | 9175695 | 9176082 | AT1G26558 intron1 | + |  |  |  |  |  |  |
| Chr1 | 9176340 | 9176759 | AT1G26558 intron3 | + |  |  |  |  |  |  |
| Chr1 | 9176838 | 9177373 | AT1G26558 intron4 | + |  |  |  |  |  |  |
| Chr1 | 9574741 | 9574865 | AT1G27565 intron1 | + |  |  |  |  |  |  |
| Chr1 | 9802501 | 9802574 | AT1G28100 intron4 | + |  |  |  |  |  |  |
| Chr1 | 9803376 | 9803488 | AT1G28100 intron8 | + |  |  |  |  |  |  |
| Chr1 | 11644266 | 11644581 | AT1G32270 intron5 | + | Chr1 | 11644394 | 11644558 | AT1TE37685 | RathE1 cons | - |
| Chr1 | 11715748 | 11715841 | AT1G32450 intron1 | - |  |  |  |  |  |  |
| Chr1 | 11716341 | 11716419 | AT1G32450 intron2 | - |  |  |  |  |  |  |
| Chr1 | 11871347 | 11871725 | AT1G32780_intron6 | - |  |  |  |  |  |  |
| Chr1 | 12650863 | 12652394 | AT1G34550 intron7 | - | Chr1 | 12651410 | 12651914 | AT1TE41310 | ATHATN3A | + |
| Chr1 | 13024455 | 13024627 | AT1G35410 intron2 | + |  |  |  |  |  |  |
| Chr1 | 13836539 | 13836822 | AT1G36622 intron1 | - |  |  |  |  |  |  |
| Chr1 | 13868086 | 13868159 | AT1G36675 intron2 | + |  |  |  |  |  |  |
| Chr1 | 13868289 | 13869168 | AT1G36675 intron3 | + |  |  |  |  |  |  |
| Chr1 | 13869226 | 13869263 | AT1G36675 intron4 | + |  |  |  |  |  |  |
| Chr1 | 17116439 | 17116811 | AT1G45191 intron3 | + | Chr1 | 17116653 | 17116793 | AT1TE56665 | ATTIRTA1 | + |
| Chr1 | 17417964 | 17418692 | AT1G47480 intron1 | + |  |  |  |  |  |  |
| Chr1 | 20023772 | 20024949 | AT1G53640 intron3 | - |  |  |  |  |  |  |
| Chr1 | 20115705 | 20115927 | AT1G53880 intron2 | + |  |  |  |  |  |  |
| Chr1 | 20273709 | 20273779 | AT1G54310 intron1 | + |  |  |  |  |  |  |
| Chr1 | 20422928 | 20423399 | AT1G54720 intron1 | + | Chr1 | 20422943 | 20423145 | AT1TE67445 | ATHILA7 | - |
| Chr1 | 20423538 | 20423623 | AT1G54720 intron2 | + |  |  |  |  |  |  |
| Chr1 | 21179305 | 21180398 | AT1G56530 intron1 | + | Chr1 | 21179997 | 21180142 | AT1TE70010 | ATHATN3 | - |
| Chr1 | 21731181 | 21733417 | AT1G58520 intron5 | + | Chr1 | 21731638 | 21732236 | AT1TE71705 | ATHATN6 | + |
| Chr1 | 21746500 | 21753883 | AT1G58602 intron1 | + | Chr1 | 21746512 | 21746739 | AT1TE71765 | ATREP10 | + |
| Chr1 | 21746500 | 21753883 | AT1G58602_intron1 | + | Chr1 | 21746740<br>21748194<br>21753390<br>21746512 | 21747327<br>21753388<br>21753542<br>21746739 | AT1TE71770<br>AT1TE71775<br>AT1TE71780<br>AT1TE71765 | ATHATN3<br>ATCOPIA8B<br>ATCOPIA67<br>ATREP10 | -<br>+<br>-<br>+ |
| Chr1 | 21809072 | 21809158 | AT1G59030 intron3 | - |  |  |  |  |  |  |
| Chr1 | 22563161 | 22563332 | AT1G61200 intron1 | - |  |  |  |  |  |  |
| Chr1 | 23510156 | 23510840 | AT1G63410 intron1 | - | Chr1 | 23510461 | 23510643 | AT1TE77415 | ATMU2 | - |
| Chr1 | 23537460 | 23537546 | AT1G63470 intron2 | - |  |  |  |  |  |  |
| Chr1 | 23537679 | 23537758 | AT1G63470 intron3 | - |  |  |  |  |  |  |
| Chr1 | 24552471 | 24555768 | AT1G65960_intron2 | + | Chr1 | 24553446<br>24553578 | 24553577<br>24554047 | AT1TE80650<br>AT1TE80655 | HELITRONY1D<br>ATREP1 | +<br>- |
| Chr1 | 24947003 | 24948885 | AT1G66880 intron1 | + | Chr1 | 24948088 | 24948353 | AT1TE81950 | RP1 AT | - |
| Chr1 | 25478154 | 25478224 | AT1G67940 intron1 | + |  |  |  |  |  |  |
| Chr1 | 25676957 | 25677311 | AT1G68470 intron1 | - |  |  |  |  |  |  |
| Chr1 | 27076371 | 27076785 | AT1G71930 intron1 | + |  |  |  |  |  |  |
| Chr1 | 27494452 | 27494548 | AT1G73110_intron1 | - |  |  |  |  |  |  |
| Chr1 | 27494642 | 27494714 | AT1G73110_intron2 | - |  |  |  |  |  |  |
| Chr1 | 27494771 | 27494842 | AT1G73110_intron3 | - |  |  |  |  |  |  |
| Chr1 | 27494950 | 27495043 | AT1G73110_intron4 | - |  |  |  |  |  |  |
| Chr1 | 27495246 | 27495342 | AT1G73110_intron5 | - |  |  |  |  |  |  |
| Chr1 | 28136009 | 28136127 | AT1G74910 intron2 | - |  |  |  |  |  |  |
| Chr1 | 28316316 | 28316841 | AT1G75450 intron2 | - |  |  |  |  |  |  |
| Chr1 | 29313005 | 29313463 | AT1G77960 intron12 | - |  |  |  |  |  |  |
| Chr1 | 29316500 | 29316609 | AT1G77980 intron7 | - |  |  |  |  |  |  |
| Chr1 | 29316643 | 29316728 | AT1G77980 intron8 | - |  |  |  |  |  |  |
| Chr1 | 29316783 | 29316882 | AT1G77980 intron9 | - |  |  |  |  |  |  |
| Chr1 | 29708673 | 29708764 | AT1G78980 intron3 | - |  |  |  |  |  |  |
| Chr1 | 29709037 | 29709107 | AT1G78980 intron4 | - |  |  |  |  |  |  |
| Chr2 | 184206 | 184467 | AT2G01422 intron1 | + |  |  |  |  |  |  |
| Chr2 | 677325 | 678063 | AT2G02520 intron1 | - |  |  |  |  |  |  |
| Chr2 | 763370 | 763468 | AT2G02720_intron1 | + |  |  |  |  |  |  |
| Chr2 | 764157 | 764267 | AT2G02720 intron2 | + |  |  |  |  |  |  |
| Chr2 | 1392364 | 1392466 | AT2G04135 intron2 | - |  |  |  |  |  |  |
| Chr2 | 1392364 | 1392466 | AT2G04135 intron2 | - |  |  |  |  |  |  |
| Chr2 | 1721900 | 1722280 | AT2G04890 intron1 | - |  |  |  |  |  |  |
| Chr2 | 1729989 | 1730092 | AT2G04925 intron1 | - |  |  |  |  |  |  |
| Chr2 | 2231679 | 2231758 | AT2G05830 intron8 | + |  |  |  |  |  |  |
| Chr2 | 2429282 | 2429365 | AT2G06210_intron1 | - |  |  |  |  |  |  |
| Chr2 | 3045668 | 3045776 | AT2G07340 intron1 | + |  |  |  |  |  |  |
| Chr2 | 3045831 | 3046041 | AT2G07340 intron2 | + |  |  |  |  |  |  |
| Chr2 | 4335212 | 4335523 | AT2G10975 intron1 | + |  |  |  |  |  |  |
| Chr2 | 4335668 | 4335726 | AT2G10975 intron2 | + |  |  |  |  |  |  |
| Chr2 | 4336397 | 4336546 | AT2G10980 intron1 | + |  |  |  |  |  |  |
| Chr2 | 5737216 | 5737428 | AT2G13770 intron1 | + |  |  |  |  |  |  |
| Chr2 | 5925824 | 5926195 | AT2G14080 intron1 | + | Chr2 | 5925853 | 5926014 | AT2TE24140 | ATLINE1A | - |
| Chr2 | 6153663 | 6153765 | AT2G14455 intron3 | + |  |  |  |  |  |  |
| Chr2 | 6214684 | 6214996 | AT2G14560 intron2 | + | Chr2 | 6214710 | 6214915 | AT2TE25290 | ATREP5 | + |
| Chr2 | 6863067 | 6863138 | AT2G15750 intron1 | + |  |  |  |  |  |  |
| Chr2 | 6863067 | 6863138 | AT2G15750 intron1 | + |  |  |  |  |  |  |
| Chr2 | 6863265 | 6863448 | AT2G15750 intron2 | + |  |  |  |  |  |  |
| Chr2 | 6863265 | 6863448 | AT2G15750 intron2 | + |  |  |  |  |  |  |
| Chr2 | 6887384 | 6887455 | AT2G15815 intron1 | - |  |  |  |  |  |  |
| Chr2 | 6887384 | 6887455 | AT2G15815 intron1 | - |  |  |  |  |  |  |
| Chr2 | 8366014 | 8366889 | AT2G19290 intron1 | - |  |  |  |  |  |  |
| Chr2 | 8569085 | 8569431 | AT2G19850 intron1 | + |  |  |  |  |  |  |
| Chr2 | 9048186 | 9048639 | AT2G21100 intron1 | - |  |  |  |  |  |  |
| Chr2 | 9344305 | 9344846 | AT2G21920 intron2 | - |  |  |  |  |  |  |
| Chr2 | 9344305 | 9344846 | AT2G21920 intron2 | - |  |  |  |  |  |  |
| Chr2 | 9652013 | 9653650 | AT2G22710 intron1 | + |  |  |  |  |  |  |
| Chr2 | 9652013 | 9653650 | AT2G22710 intron1 | + |  |  |  |  |  |  |
| Chr2 | 10289490 | 10289580 | AT2G24200 intron10 | - |  |  |  |  |  |  |
| Chr2 | 11123577 | 11123649 | AT2G26120 intron1 | + |  |  |  |  |  |  |
| Chr2 | 11167633 | 11167724 | AT2G26240 intron1 | - |  |  |  |  |  |  |
| Chr2 | 13020840 | 13020991 | AT2G30575 intron1 | - |  |  |  |  |  |  |
| Chr2 | 13094419 | 13094557 | AT2G30730 intron5 | + |  |  |  |  |  |  |
| Chr2 | 15473236 | 15473914 | AT2G36870 intron1 | - |  |  |  |  |  |  |
| Chr2 | 15681944 | 15682024 | AT2G37370 intron14 | + |  |  |  |  |  |  |
| Chr2 | 16022527 | 16023705 | AT2G38255 intron2 | - | Chr2 | 16023403 | 16023576 | AT2TE71735 | ARNOLDY2 | + |
| Chr2 | 16333809 | 16334129 | AT2G39160 intron1 | - |  |  |  |  |  |  |
| Chr2 | 16889584 | 16890860 | AT2G40440 intron2 | - |  |  |  |  |  |  |
| Chr2 | 17362319 | 17362890 | AT2G41640 intron3 | + | Chr2 | 17362655 | 17362821 | AT2TE78320 | ATSINE2A | + |
| Chr2 | 17363952 | 17364032 | AT2G41650 intron1 | + |  |  |  |  |  |  |
| Chr2 | 18910778 | 18911002 | AT2G45960 intron1 | + |  |  |  |  |  |  |
| Chr2 | 18911520 | 18911607 | AT2G45960 intron3 | + |  |  |  |  |  |  |
| Chr3 | 54929 | 55008 | AT3G01160 intron1 | - |  |  |  |  |  |  |
| Chr3 | 1555396 | 1561143 | AT3G05410 intron2 | + | Chr3 | 1555420 | 1560680 | AT3TE06550 | ATLINE2 | - |
| Chr3 | 1674500 | 1674854 | AT3G05685 intron3 | - |  |  |  |  |  |  |
| Chr3 | 2264181 | 2264272 | AT3G07150 intron2 | - |  |  |  |  |  |  |
| Chr3 | 2429653 | 2431126 | AT3G07610 intron8 | + |  |  |  |  |  |  |
| Chr3 | 2557895 | 2557995 | AT3G08020 intron1 | - |  |  |  |  |  |  |
| Chr3 | 2558477 | 2558619 | AT3G08020 intron2 | - |  |  |  |  |  |  |
| Chr3 | 2600851 | 2601680 | AT3G08560 intron3 | + | Chr3 | 2600864 | 2601601 | AT3TE10950 | TAG3N1 | - |
| Chr3 | 3424305 | 3424419 | AT3G10950 intron3 | + |  |  |  |  |  |  |
| Chr3 | 3498332 | 3498432 | AT3G11165 intron3 | - |  |  |  |  |  |  |
| Chr3 | 5861606 | 5861997 | AT3G17185 intron1 | + |  |  |  |  |  |  |
| Chr3 | 6705625 | 6705716 | AT3G19350 intron1 | + |  |  |  |  |  |  |
| Chr3 | 6930151 | 6930274 | AT3G19920 intron1 | - |  |  |  |  |  |  |
| Chr3 | 7704174 | 7704427 | AT3G21870 intron1 | - |  |  |  |  |  |  |
| Chr3 | 8615772 | 8616255 | AT3G23850 intron2 | + |  |  |  |  |  |  |
| Chr3 | 8977602 | 8977756 | AT3G24610 intron4 | - |  |  |  |  |  |  |
| Chr3 | 8977844 | 8978051 | AT3G24610 intron5 | - |  |  |  |  |  |  |
| Chr3 | 9787963 | 9788104 | AT3G26616 intron1 | - |  |  |  |  |  |  |
| Chr3 | 9977744 | 9977817 | AT3G27040 intron5 | - |  |  |  |  |  |  |
| Chr3 | 10224365 | 10224427 | AT3G27600 intron2 | - |  |  |  |  |  |  |
| Chr3 | 10224550 | 10224572 | AT3G27600 intron3 | - |  |  |  |  |  |  |
| Chr3 | 10449512 | 10450226 | AT3G28070 intron4 | + | Chr3 | 10449619 | 10450104 | AT3TE43415 | ATHATN1 | - |
| Chr3 | 10976485 | 10976733 | AT3G28950 intron1 | - |  |  |  |  |  |  |
| Chr3 | 11103432 | 11103550 | AT3G29130 intron4 | - |  |  |  |  |  |  |
| Chr3 | 14082962 | 14083054 | AT3G33528 intron1 | - |  |  |  |  |  |  |
| Chr3 | 14083235 | 14083352 | AT3G33528 intron2 | - |  |  |  |  |  |  |
| Chr3 | 14083480 | 14083901 | AT3G33528 intron3 | - |  |  |  |  |  |  |
| Chr3 | 14083985 | 14084138 | AT3G33528 intron4 | - |  |  |  |  |  |  |
| Chr3 | 14689163 | 14689999 | AT3G42570_intron1 | + | Chr3 | 14689254<br>14689700 | 14689695<br>14689877 | AT3TE60115<br>AT3TE60120 | BRODYAGA1A<br>HELITRONY1B | -<br>+ |
| Chr3 | 14690857 | 14691128 | AT3G42580 intron1 | + |  |  |  |  |  |  |
| Chr3 | 14690857 | 14691128 | AT3G42580 intron1 | + |  |  |  |  |  |  |
| Chr3 | 14924398 | 14924552 | AT3G42820 intron13 | - |  |  |  |  |  |  |
| Chr3 | 14924398 | 14924552 | AT3G42820 intron13 | - |  |  |  |  |  |  |
| Chr3 | 15221585 | 15221675 | AT3G43260 intron5 | + |  |  |  |  |  |  |
| Chr3 | 16010313 | 16010428 | AT3G44330 intron12 | + |  |  |  |  |  |  |
| Chr3 | 16010519 | 16010608 | AT3G44330 intron13 | + |  |  |  |  |  |  |
| Chr3 | 16071075 | 16071124 | AT3G44440 intron1 | + |  |  |  |  |  |  |
| Chr3 | 16302318 | 16302404 | AT3G44755 intron2 | + |  |  |  |  |  |  |
| Chr3 | 16527567 | 16527652 | AT3G45140 intron4 | + |  |  |  |  |  |  |
| Chr3 | 16816916 | 16818543 | AT3G45780 intron1 | + | Chr3 | 16817888 | 16817997 | AT3TE68105 | RathE2 cons | - |
| Chr3 | 16840352 | 16840835 | AT3G45820 intron2 | - |  |  |  |  |  |  |
| Chr3 | 17059758 | 17059835 | AT3G46380 intron2 | - |  |  |  |  |  |  |
| Chr3 | 17104524 | 17104597 | AT3G46480 intron5 | + |  |  |  |  |  |  |
| Chr3 | 17491557 | 17491789 | AT3G47460 intron14 | + |  |  |  |  |  |  |
| Chr3 | 17491975 | 17492142 | AT3G47460 intron15 | + |  |  |  |  |  |  |
| Chr3 | 18254533 | 18254849 | AT3G49230 intron1 | + |  |  |  |  |  |  |
| Chr3 | 18270722 | 18270798 | AT3G49280 intron3 | + |  |  |  |  |  |  |
| Chr3 | 19533208 | 19538341 | AT3G52700_intron3 | - | Chr3 | 19533230<br>19533925<br>19536332<br>19536457 | 19533871<br>19534234<br>19536456<br>19537958 | AT3TE79290<br>AT3TE79300<br>AT3TE79315<br>AT3TE79320 | HELITRONY1D<br>ATCOPIA49<br>ARNOLDY2<br>HELITRONY1E | +<br>-<br>-<br>- |
| Chr3 | 22525701 | 22525896 | AT3G60950 intron4 | + |  |  |  |  |  |  |
| Chr3 | 22525924 | 22526052 | AT3G60950 intron5 | + |  |  |  |  |  |  |
| Chr4 | 667503 | 668111 | AT4G01533 intron1 | - |  |  |  |  |  |  |
| Chr4 | 668235 | 668346 | AT4G01533 intron2 | - |  |  |  |  |  |  |
| Chr4 | 1288833 | 1289330 | AT4G02910 intron2 | - |  |  |  |  |  |  |
| Chr4 | 1447426 | 1447928 | AT4G03300 intron1 | + |  |  |  |  |  |  |
| Chr4 | 1447426 | 1447928 | AT4G03300 intron1 | + |  |  |  |  |  |  |
| Chr4 | 1448678 | 1449451 | AT4G03300 intron3 | + |  |  |  |  |  |  |
| Chr4 | 2035795 | 2036278 | AT4G04221_intron2 | - |  |  |  |  |  |  |
| Chr4 | 2036818 | 2036891 | AT4G04223_intron1 | + |  |  |  |  |  |  |
| Chr4 | 2036934 | 2037600 | AT4G04223_intron2 | + |  |  |  |  |  |  |
| Chr4 | 2293863 | 2293970 | AT4G04580 intron1 | - |  |  |  |  |  |  |
| Chr4 | 2294234 | 2294382 | AT4G04580 intron3 | - |  |  |  |  |  |  |
| Chr4 | 2577328 | 2577780 | AT4G05030 intron2 | - |  |  |  |  |  |  |
| Chr4 | 2577849 | 2577934 | AT4G05030 intron3 | - |  |  |  |  |  |  |
| Chr4 | 5230561 | 5230963 | AT4G08280 intron1 | + | Chr4 | 5230692 | 5230870 | AT4TE21780 | ATREP10A | + |
| Chr4 | 5231054 | 5231121 | AT4G08280 intron2 | + |  |  |  |  |  |  |
| Chr4 | 5231245 | 5231853 | AT4G08280 intron3 | + |  |  |  |  |  |  |
| Chr4 | 5643999 | 5644447 | AT4G08860 intron1 | + |  |  |  |  |  |  |
| Chr4 | 5643999 | 5644447 | AT4G08860 intron1 | + |  |  |  |  |  |  |
| Chr4 | 5782197 | 5783226 | AT4G09012 intron3 | - | Chr4 | 5782384 | 5782761 | AT4TE24280 | ATTIRX1B | - |
| Chr4 | 6016029 | 6016093 | AT4G09490 intron1 | - |  |  |  |  |  |  |
| Chr4 | 7799731 | 7801215 | AT4G13420_intron6 | - | Chr4 | 7799736<br>7799859 | 7799858<br>7800628 | AT4TE34210<br>AT4TE34215 | HELITRON4<br>BRODYAGA2 | +<br>- |
| Chr4 | 7801439 | 7801523 | AT4G13420 intron7 | - |  |  |  |  |  |  |
| Chr4 | 7801608 | 7802102 | AT4G13420 intron8 | - |  |  |  |  |  |  |
| Chr4 | 8147748 | 8148098 | AT4G14140 intron1 | + |  |  |  |  |  |  |
| Chr4 | 9490379 | 9490478 | AT4G16860 intron3 | - |  |  |  |  |  |  |
| Chr4 | 9490812 | 9490994 | AT4G16860 intron4 | - |  |  |  |  |  |  |
| Chr4 | 9522091 | 9522239 | AT4G16920 intron5 | - |  |  |  |  |  |  |
| Chr4 | 9522534 | 9524001 | AT4G16920 intron6 | - | Chr4 | 9522668 | 9523966 | AT4TE43010 | HELITRONY1A | - |
| Chr4 | 9541555 | 9541670 | AT4G16950 intron4 | - |  |  |  |  |  |  |
| Chr4 | 10050386 | 10050738 | AT4G18150 intron1 | + |  |  |  |  |  |  |
| Chr4 | 10897979 | 10898448 | AT4G20170 intron1 | + |  |  |  |  |  |  |
| Chr4 | 11240907 | 11241765 | AT4G21060 intron1 | + |  |  |  |  |  |  |
| Chr4 | 11447636 | 11447725 | AT4G21520 intron1 | - |  |  |  |  |  |  |
| Chr4 | 12021982 | 12022063 | AT4G22940 intron1 | - |  |  |  |  |  |  |
| Chr4 | 12022292 | 12022374 | AT4G22940 intron2 | - |  |  |  |  |  |  |
| Chr4 | 12022693 | 12022780 | AT4G22940 intron3 | - |  |  |  |  |  |  |
| Chr4 | 12293960 | 12294034 | AT4G23560 intron1 | - |  |  |  |  |  |  |
| Chr4 | 12750367 | 12752787 | AT4G24730 intron1 | - | Chr4 | 12751593 | 12752065 | AT4TE59555 | ATCOPIA7 | + |
| Chr4 | 13038629 | 13039292 | AT4G25530 intron2 | + |  |  |  |  |  |  |
| Chr4 | 13472651 | 13472754 | AT4G26730 intron1 | + |  |  |  |  |  |  |
| Chr4 | 13549218 | 13549334 | AT4G26980 intron1 | - |  |  |  |  |  |  |
| Chr4 | 13549753 | 13550014 | AT4G26980 intron3 | - |  |  |  |  |  |  |
| Chr4 | 16177133 | 16177205 | AT4G33700 intron3 | - |  |  |  |  |  |  |
| Chr4 | 16177280 | 16177405 | AT4G33700 intron4 | - |  |  |  |  |  |  |
| Chr4 | 16325873 | 16325984 | AT4G34071 intron1 | + |  |  |  |  |  |  |
| Chr4 | 16326169 | 16326622 | AT4G34071 intron2 | + |  |  |  |  |  |  |
| Chr4 | 16811564 | 16811659 | AT4G35350 intron3 | + |  |  |  |  |  |  |
| Chr5 | 2107401 | 2107483 | AT5G06805 intron2 | - |  |  |  |  |  |  |
| Chr5 | 2579459 | 2579614 | AT5G08055 intron1 | + |  |  |  |  |  |  |
| Chr5 | 2779399 | 2779487 | AT5G08570 intron3 | + |  |  |  |  |  |  |
| Chr5 | 3253261 | 3253846 | AT5G10340 intron1 | + | Chr5 | 3253482 | 3253614 | AT5TE11835 | VANDAL18NA | + |
| Chr5 | 3372737 | 3372831 | AT5G10670 intron1 | - |  |  |  |  |  |  |
| Chr5 | 5008400 | 5008933 | AT5G15420 intron1 | - |  |  |  |  |  |  |
| Chr5 | 6741017 | 6741135 | AT5G19950 intron1 | - |  |  |  |  |  |  |
| Chr5 | 7631658 | 7631755 | AT5G22840 intron2 | - |  |  |  |  |  |  |
| Chr5 | 8162790 | 8163154 | AT5G24130 intron2 | + |  |  |  |  |  |  |
| Chr5 | 8233223 | 8233960 | AT5G24240 intron2 | - |  |  |  |  |  |  |
| Chr5 | 8234305 | 8234580 | AT5G24250 intron1 | - |  |  |  |  |  |  |
| Chr5 | 8622552 | 8622625 | AT5G25040 intron1 | + |  |  |  |  |  |  |
| Chr5 | 8622886 | 8622961 | AT5G25040 intron2 | + |  |  |  |  |  |  |
| Chr5 | 8623297 | 8623371 | AT5G25040 intron4 | + |  |  |  |  |  |  |
| Chr5 | 8623971 | 8624110 | AT5G25040 intron6 | + |  |  |  |  |  |  |
| Chr5 | 10786467 | 10786787 | AT5G28760 intron1 | + |  |  |  |  |  |  |
| Chr5 | 12272534 | 12272928 | AT5G32621 intron1 | + |  |  |  |  |  |  |
| Chr5 | 13111250 | 13111382 | AT5G34850 intron8 | - |  |  |  |  |  |  |
| Chr5 | 13934502 | 13934642 | AT5G35760 intron1 | + |  |  |  |  |  |  |
| Chr5 | 14067414 | 14067506 | AT5G35930 intron1 | - |  |  |  |  |  |  |
| Chr5 | 15155341 | 15155817 | AT5G38005 intron1 | + |  |  |  |  |  |  |
| Chr5 | 20978824 | 20979207 | AT5G51650 intron2 | - |  |  |  |  |  |  |
| Chr5 | 21161354 | 21162095 | AT5G52070 intron1 | + |  |  |  |  |  |  |
| Chr5 | 21162209 | 21163578 | AT5G52070 intron2 | + | Chr5 | 21162772 | 21163127 | AT5TE76210 | ATCOPIA68 | - |
| Chr5 | 21164738 | 21164823 | AT5G52070 intron7 | + |  |  |  |  |  |  |
| Chr5 | 21170308 | 21170395 | AT5G52100 intron1 | + |  |  |  |  |  |  |
| Chr5 | 21170519 | 21170606 | AT5G52100 intron2 | + |  |  |  |  |  |  |
| Chr5 | 21170835 | 21170920 | AT5G52100 intron4 | + |  |  |  |  |  |  |
| Chr5 | 21171017 | 21171157 | AT5G52100 intron5 | + |  |  |  |  |  |  |
| Chr5 | 21759465 | 21759616 | AT5G53560 intron1 | + |  |  |  |  |  |  |
| Chr5 | 21759715 | 21759947 | AT5G53560 intron2 | + |  |  |  |  |  |  |
| Chr5 | 21760015 | 21760102 | AT5G53560 intron3 | + |  |  |  |  |  |  |
| Chr5 | 22407943 | 22409325 | AT5G55250 intron1 | - |  |  |  |  |  |  |
| Chr5 | 23361462 | 23361538 | AT5G57670 intron3 | - |  |  |  |  |  |  |
| Chr5 | 23361697 | 23361809 | AT5G57670 intron4 | - |  |  |  |  |  |  |

